# Host metabolic pathways essential for malaria and related hemoparasites in the infection of nucleated cells

**DOI:** 10.1101/2023.09.27.559824

**Authors:** Marina Maurizio, Maria Masid, Kerry Woods, Reto Caldelari, John G. Doench, Arunasalam Naguleswaran, Denis Joly, Martín González Fernández, Jonas Zemp, Mélanie Borteele, Vassily Hatzimanikatis, Volker Heussler, Sven Rottenberg, Philipp Olias

## Abstract

Apicomplexan parasite diseases, including malaria (*Plasmodium*) and theileriosis (*Theileria*), pose a significant threat to global health and the socioeconomic well-being of low-income countries. Despite recent advances, the common host metabolic proteins essential for these highly auxotrophic pathogens remain elusive. Here, we present a comprehensive investigation integrating a metabolic model of *P. falciparum* parasites in hepatocytes and a genome-wide CRISPR screen targeting *Theileria* schizont-infected macrophages. We reveal unifying host metabolic enzymes critical for the intracellular survival of these related hematozoa. We show that pathways such as host purine and heme biosynthesis are essential for both *Theileria* survival and *Plasmodium* liver development, while genes involved in glutathione and polyamine biosynthesis are predicted to be essential for *Plasmodium* only under certain metabolic conditions. Our work highlights the importance of host porphyrins for the viability of liver-stage *Plasmodium*. Shared parasite vulnerabilities provide a resource for exploring alternative therapeutic approaches to combat these crippling diseases.

## INTRODUCTION

Apicomplexa are a large group of unicellular parasites, with more than 6,000 described species and an estimated million(s) yet to be identified.^1^ The obligate intracellular organisms can cause severe and life-threatening diseases in humans and animals, of which malaria and theileriosis are particularly devastating. Malaria-causing *Plasmodium* is the most impactful human parasite, with over 245 million malaria cases and more than 600,000 deaths reported annually.^2^ Another socio-economically important apicomplexan is *Theileria*, which infects cattle and causes fatal tropical theileriosis (*T. annulata*) in the Middle East and Asia, and east coast fever (*T. parva*) in sub-Saharan Africa.^3–6^ *Theileria* parasites have the unique ability among eukaryotic pathogens to induce cancer-like transformation of invaded cells, characterized by sustained proliferation, invasiveness, and altered metabolism.^7,8^

A hallmark of all apicomplexans is that they are auxotrophic for multiple metabolites that need to be taken up from the extra-parasitic environment. This requires complex interactions with the host cell and manipulation of key biological processes.^9–11^ Our knowledge of the host metabolic proteins that are essential for the efficient development of these protozoa in their intracellular niche is still very limited. We hypothesized that apicomplexans belonging to different genera rely on shared host metabolic proteins for their development and intracellular survival, and that direct targeting of the host cell and nutrient uptake mechanisms could provide opportunities for new drug development or repurposing. To investigate whether there is an overlap in key host metabolic pathways, we focused on what are arguably the two most important genera of hemoparasites, *Plasmodium* and *Theileria*. A common feature of these two parasites is that, unlike other hematozoa (such as related *Babesia*), they undergo a huge expansion within nucleated cells termed schizogony prior to their red blood cell stage. This occurs within hepatocytes (*Plasmodium*) and white blood cells (*Theileria*). Following this expansion, the parasites develop into large numbers of merozoites that subsequently infect red blood cells.

Reverse genetic screens, using tools such as CRISPR/Cas9 and RNAi, have proven to be highly effective in the identification of fitness-conferring genes for intracellular pathogens. Large-scale CRISPR screens have been applied to study gene function in a wide variety of living systems, including mammalian cells,^12^ bacteria,^13^ and apicomplexan parasites.^14–16^ Only a few large-scale screens have focused on parasite-specific host gene essentiality, including the murine malaria parasite *P. yoelli*, and the zoonotic parasites *Cryptosporidium parvum* and *Toxoplasma gondii*.^17–19^ Studies of *P. falciparum* and *Theileria*-infected host cells are still lacking. While *Theileria* schizonts can be studied in cell culture with quantitative proteomics and host genomic technologies, *Plasmodium* schizont infected hepatocytes are technically difficult to study in large numbers, and large-scale host metabolic data on the major malaria causing species, *P. falciparum*, are lacking.

Genome-scale metabolic models (GEMs) have emerged as invaluable tools in computational systems biology, providing a comprehensive and systems-level understanding of cellular metabolism.^20^ By integrating genomic, biochemical, and physiological data, these models provide a holistic representation of the cellular metabolic networks.^21^ To date, several GEMs have been developed for apicomplexan parasites including *Plasmodium*.^22–24^ Advances in genome annotation, enzyme localization, and definition of available substrates have allowed for a more fine-tuned prediction of essential properties and metabolic activities within *Plasmodium* in recent years.^16,25^ One of the most powerful applications of GEMs is their ability to investigate gene essentiality for cell survival and function. By simulating the metabolic fluxes and interactions within a cell, the effects of genetic perturbations on cell growth can be systematically analyzed, advancing our fundamental understanding of cellular processes.^26, 27^

In this study, we combined computational and experimental approaches to investigate the intricate metabolic interactions between apicomplexan parasites and their nucleated host cells. We reconstructed metabolic models to investigate the dependency of the *P. falciparum* liver stages on host metabolic genes. In parallel, we performed a CRISPR/Cas9 drop-out screen using a novel bovine genome-wide library in *T. annulata* schizont-infected macrophages, the life cycle stage most comparable to the *P. falciparum* liver stage. We identified a set of fitness-conferring genes that are required for the survival of infected macrophages but are dispensable for uninfected macrophages. Our results suggest that multiple host cell biosynthesis pathways and specific metabolic proteins are essential for both *Plasmodium* and *Theileria* parasites. Specifically, we found that *Plasmodium* schizonts depend on the host heme biosynthesis pathway to efficiently develop within the liver. These findings provide new insights into the metabolic interactions between apicomplexan parasites and their host cell environment that could be exploited to develop new strategies for the treatment of disease.

## RESULTS

### Modeling host-parasite metabolic interactions

To study the metabolic interactions between the malaria parasite *P. falciparum* and the human hepatocyte, we developed a systems biology approach to reconstruct a metabolic model that integrates information on the metabolism of both the parasite and the specific host cell, and how the parasite scavenges nutrients from the host. We first reconstructed a human hepatocyte-specific metabolic model (**Figure 1A**) by collecting information from the human protein atlas (www.proteinatlas.org)^28^ and previous hepatocyte metabolic networks.^29,30^ We then used this information to extract all reactions associated with genes expressed in liver cells from the thermodynamically curated human genome-scale model Recon 3D.^31,32^ Next, we built a liver-specific *P. falciparum* metabolic model (liver-iPfa) from the genome-scale model iPfa^25^ by defining as nutrients for the parasite those metabolites present in the cytosol of the reconstructed hepatocyte model (**Figure 1B**). Subsequently, the liver-iPfa model was used to study the alternative nutritional requirements of the parasite and the utilization of its available biochemical pathways for biomass synthesis and growth. In addition, we introduce the concept of the *parasitosome*, a reaction that summarizes the metabolic interactions of an intracellular parasite within its host cell. The *parasitosome* reaction is formulated by first identifying the metabolic reactions that the parasite needs for growth under given conditions inside the host cell. These reactions are then combined into one single reaction where substrates represent nutrients that the parasite takes up from the host’s cytosol, and products represent metabolites that the parasite secretes into the host. Given the biochemistry of the host and the flexibility of the parasite’s metabolism to adapt to varying environmental conditions, alternative *parasitosomes* may exist, representing different metabolic states of the parasites (**Figure S1A**). The alternative *parasitosomes* are then integrated into the hepatocyte model creating a host-parasite model that allows us to study the dependency of the *P. falciparum* schizont on the hepatocyte’s metabolism (**Figure 1C**). This approach allows the analysis of possible scenarios of nutrient availability to *P. falciparum* once it has infected the hepatocyte, the metabolism used by the parasite to synthesize biomass building blocks under the specific nutritional conditions, and the essentiality of host genes for parasite survival given the different physiological conditions.

**Figure 1.**
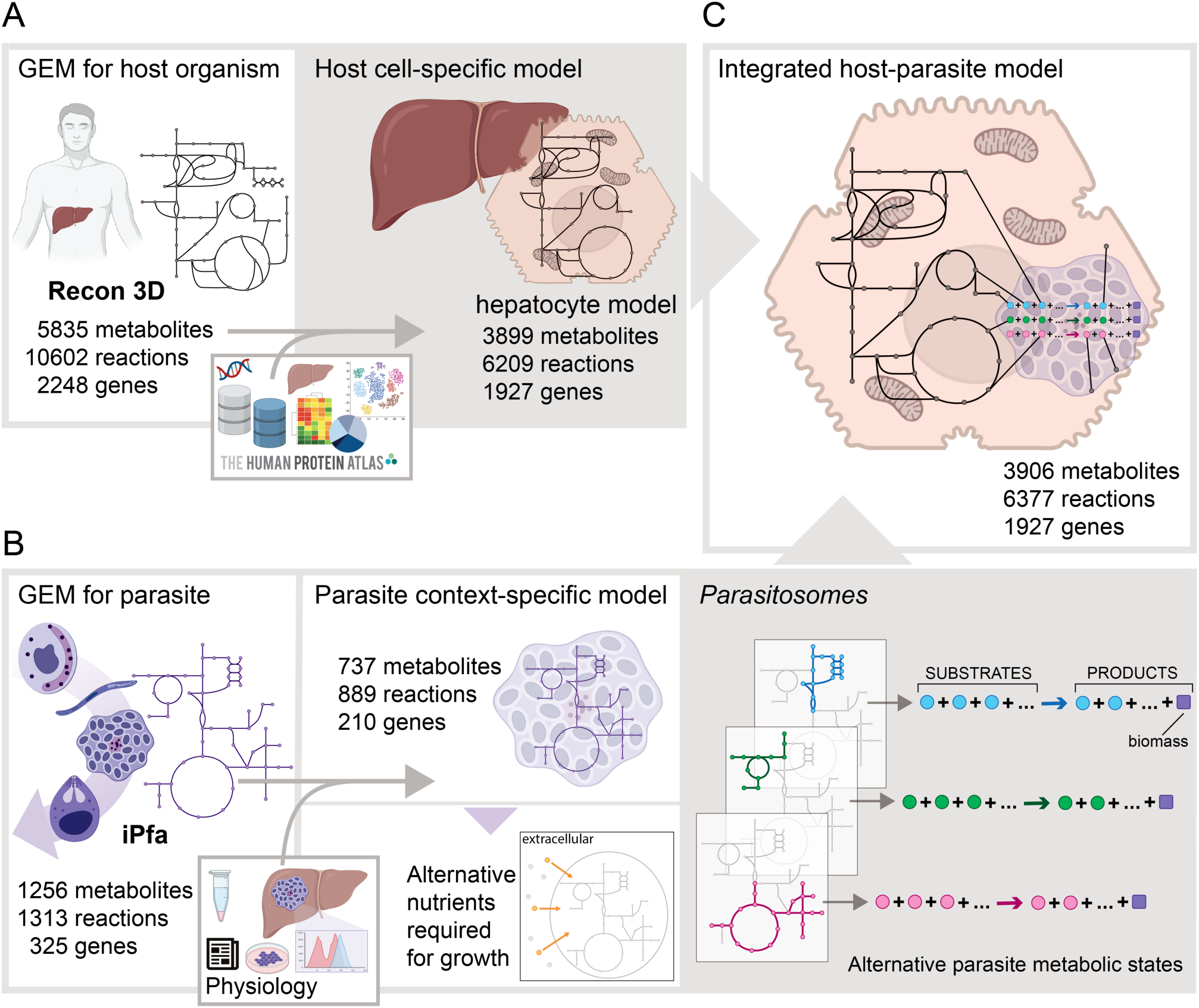
Systems biology workflow to study host-parasite metabolic interactions of *Plasmodium falciparum* in the liver. (**A**) Reconstruction of a hepatocyte-specific model from the human genome-scale metabolic model by integrating omics data. (**B**) Reconstruction of a liver-specific *P. falciparum* metabolic model from the iPfa genome-scale model by restricting the extra-parasite environment to metabolites available in the cytosol of the liver metabolic model. The reconstructed liver-iPfa model is used to analyze the nutritional requirements of the parasites and to formulate *parasitosome* reactions representing different metabolic states that the parasites can acquire to adapt to their environment and synthesize biomass. (**C**) Reconstruction of an integrated host-parasite model, by incorporating the *parasitosome* reactions into the liver metabolic model.

### Host-parasite metabolic interactions and adaptation of *P. falciparum*’s metabolism to available nutrients

We used our model to investigate two scenarios: a simulation of a parasite in a rich extra-parasitic environment inside the hepatocyte, with access to 165 nutrients (**Figure 2A**, **Table S1** and methods details), and a parasite in a compromised extra-parasitic environment inside the hepatocyte, when only a minimal set of 47 nutrients is available (**Figure 2A**, **S1B**). We then examined how the parasite utilizes the available nutrients and its own biochemistry to grow in both cases. We found that *P. falciparum* schizonts in a compromised extra-parasitic environment need to engage in more intracellular reactions than parasites that are in a rich environment (**Figures 2B**, **S2C**). Based on the computational results, when the parasites are in a rich nutritional environment, they take up a greater number of nutrients, including amino acids, saccharides, fatty acids, and nucleotides (**Figures 2C**, **S2D**). However, when *P. falciparum* finds a compromised environment inside the hepatocyte, the parasites will have to synthesize unavailable metabolites, relying on their own metabolism and engaging intracellular reactions from central carbon pathways, fatty acid metabolism, pyrimidine metabolism and vitamin metabolism (**Figure 2B**), while elevated levels of by-products such as fatty acids and detoxification molecules might be secreted into the hepatocyte’s cytosol (**Figure 2C**).

**Figure 2.**
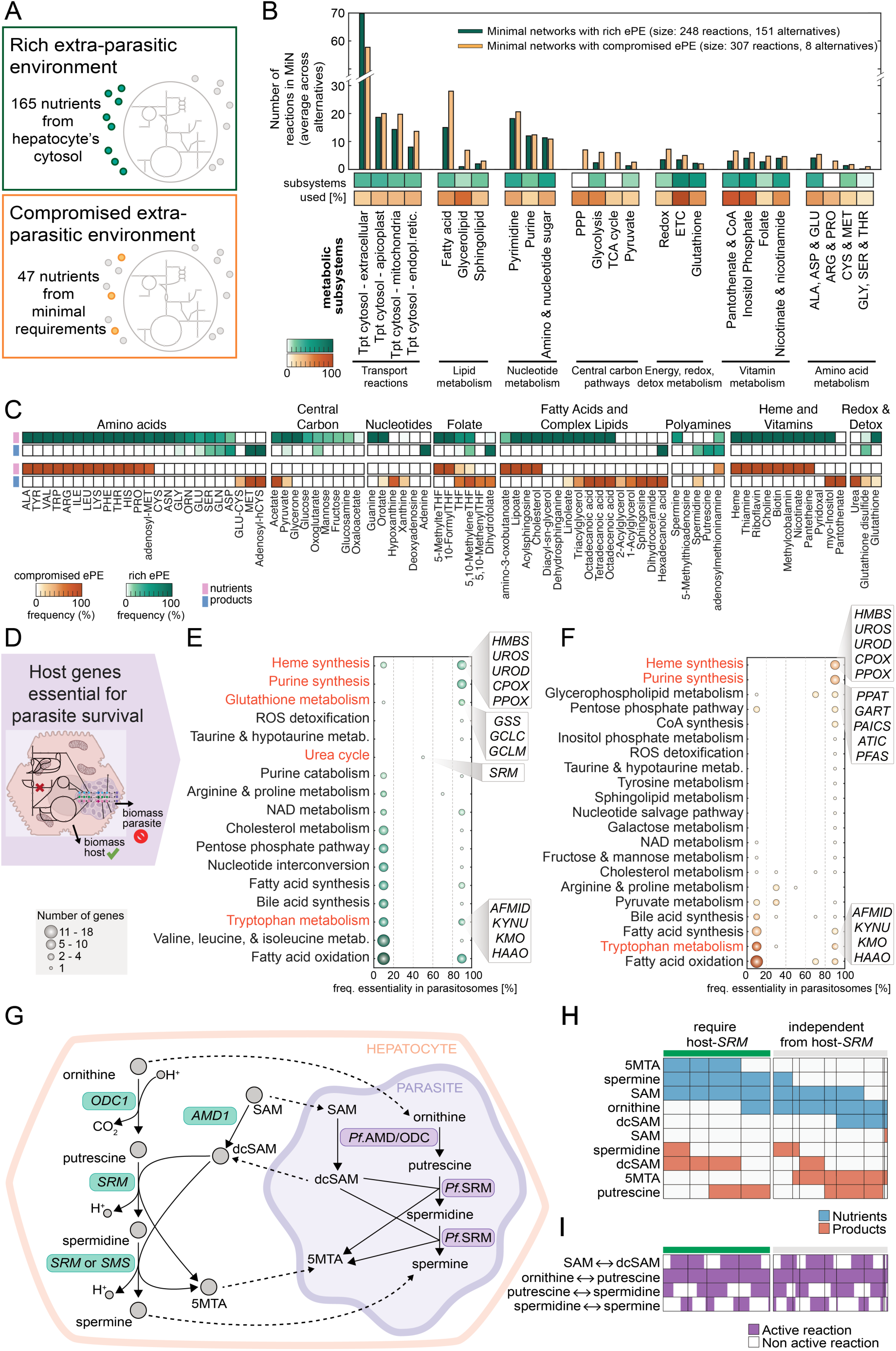
Analysis of *P. falciparum* metabolic states and host genes essential for parasite growth. (**A**) Case study I – rich extra-parasitic environment: *P. falciparum* has access to the entire hepatocyte’s cytosol metabolome. Case study II – compromised extra-parasitic environment: *P. falciparum* has access to a minimal set of nutrients from the hepatocyte’s cytosol. (**B**) Assignment to metabolic subsystems of the reactions that compose the minimal networks for the different *parasitosomes*, when considering a rich (green) or a compromised (orange) extra-parasitic environment (ePE), in terms of number of reactions and percentage of subsystem coverage for selected pathways. (**C**) Metabolic composition of the alternative *parasitosomes* in case study I (rich ePE, green) and II (compromised ePE, orange). *Parasitosomes* are representations of the nutrients taken up from and products secreted to the hepatocyte by parasites in different metabolic states. (**D**) Scheme of gene essentiality analysis performed in the integrated host-parasite model (**E-F**) Hepatocyte genes essential for *P. falciparum* in case study I (rich ePE, green) and in case study II (compromised ePE, orange) for selected pathways. Classification by associated metabolic processes, number of genes, and frequency of essentiality across *parasitosomes*. (**G**) Host-parasite interactions for polyamine synthesis. (**H**) Nutrients used and products secreted by the parasites into the hepatocyte for the polyamine’s pathway. Separated into parasites that require the hepatocyte’s *SRM* gene for survival (green) and those that survive independently of the host’s *SRM* gene (grey). Column divisions group *parasitosomes* with the same metabolic flux profile for the polyamine synthesis (**I**) Activity of the polyamine pathway in *parasitosomes* in relation to their dependence on the host’s SRM gene. SAM: adenosylmethionine, dcSAM: decarboxylated SAM, 5MTA: 5-methylthioadenosine.

Host-directed therapies in host-parasite infections are of increasing scientific interest, as they offer promising therapeutic strategies to disrupt the parasite’s life cycle, develop effective treatments, and mitigate drug resistance.^33,34^ Therefore, we used the reconstructed models to investigate which hepatocyte metabolic genes are essential for *P. falciparum* survival but dispensable for hepatocyte growth. To this end, we performed gene essentiality analyses in the healthy hepatocyte model and in the hepatocyte-*P. falciparum* integrated model (**Figure 2D**). By *in silico* knocking out of each of the 1927 metabolic genes in the hepatocyte model, we computationally identified 55 essential genes for the healthy hepatocyte (**Figure S3A**). We then simulated the knockout of each gene in the hepatocyte-parasite model to investigate the effect on parasite survival. In the case of parasites growing in a rich extra-parasitic environment, 209 hepatocyte genes from 76 metabolic pathways are essential for at least one *parasitosome*, and not for the hepatocyte. Essential pathways include heme synthesis, purine synthesis, glutathione metabolism, urea cycle, tryptophan metabolism, and fatty acid oxidation (**Figure 2E**, **S2A**). In the case of *P. falciparum* growing in a compromised extra-parasitic environment inside the hepatocyte, we identified 110 genes that are essential for at least one *parasitosome* and dispensable for the hepatocyte, belonging to 57 pathways including heme synthesis, purine metabolism, pentose phosphate pathway, and fatty acid metabolism (**Figure 2F**, **S2B**).

Interestingly, the availability of nutrients inside the hepatocyte and the metabolic state of the parasite (different *parasitosomes*) result in varying gene essentiality (**Figure S3B-C**), highlighting the flexible way *P. falciparum* can interact with the hepatocyte. This adaptability might allow the parasite to develop drug resistance and is therefore an important factor to consider when looking for drug targets. For example, when *P. falciparum* has access to a rich nutrient environment in the hepatocyte, it relies on the uptake of these nutrients to synthesize biomass precursors, and so host genes of a particular pathway will be essential. This is the case for the host glutathione synthesis genes *GSS*, *GCLC* and *GCLM*, which provide glutathione to the *parasitosomes*. However, *P. falciparum* has its own glutathione synthesis, which is active when glutathione is scarce, as in a compromised extra-parasitic environment of the host cell. Another example is the polyamine synthesis gene *SRM*, which is only essential for 50% of the *parasitosomes*, suggesting that this gene is essential under specific metabolic states of *P. falciparum*. The *parasitosome* approach allowed us to investigate further which metabolic states of the parasite lead to a specific essentiality. To illustrate this, we examined the interactions between hepatocytes and parasites for polyamines (**Figure 2G**). Our results show that *SRM* is essential when *P. falciparum* preferentially takes up spermine, adenosylmethionine (SAM) and 5-methylthioadenosine (5MTA) from the hepatocyte and secretes decarboxylated adenosylmethionine (dcSAM), depending on the hepatocyte polyamine synthesis to produce spermine and clear the parasite by-products (**Figure 2H-I**). However, *P. falciparum* can synthesize spermine from ornithine using its own polyamine synthesis, in which case *SRM* is not essential because the *parasitosomes* preferentially transport ornithine from the hepatocyte, synthesize spermine, and secrete 5MTA into the hepatocyte (**Figure 2H-I**).

We identified 19 genes that are essential for all *parasitosomes*, meaning that, based on the computational model, these genes are necessary for the survival of *P. falciparum* under all metabolic conditions. Among them, we found genes from the heme synthesis pathway (*HMBS, UROS, UROD, CPOX* and *PPOX*), purine synthesis pathway (*PPAT, GART, PAICS, ATIC,* and *PFAS*) and tryptophan metabolism (*AFMID, KYNU, KMO,* and *HAAO*) (**Figure S3D**). The other genes found are involved in NAD metabolism (*QPRT*), taurine and hypotaurine metabolism (*CDO1*), pentose phosphate pathway (*RPIA*), extracellular transport (*SLC11A2*), and cholesterol metabolism (*SOAT1*).

### Genome-wide CRISPR/Cas9 screen in *Theileria annulata* schizont infected host cells identifies parasite-specific essential host genes

Although *Plasmodium* parasites are among the most studied pathogens,^35^ large-scale experimental studies of the liver stage are underrepresented. *Theileria* parasites also develop into a multinucleated schizont in bovine leukocytes, and this life cycle stage shares many similarities with the developmental *Plasmodium* liver stage. Therefore, we hypothesized that both hematozoans share core host gene dependencies and rely on the corresponding cellular pathways for successful intracellular development. CRISPR/Cas9 loss-of-function screens are among the most valuable techniques for these investigations, and the hyperproliferative behavior of *Theileria*-infected cells makes them suitable for large-scale genetic screens. We therefore performed a genome-wide CRISPR/Cas9 dropout screen in *T. annulata*-infected bovine macrophages (TaC12; **Figure 3A**). For this purpose, we generated a CRISPR sgRNA library based on the bovine ARS-UCD1.2 genome (published in 2018) containing, on average, 4 guides per gene, for a total library size of ∼80,000 sgRNAs (**Figure 3B**).

**Figure 3.**
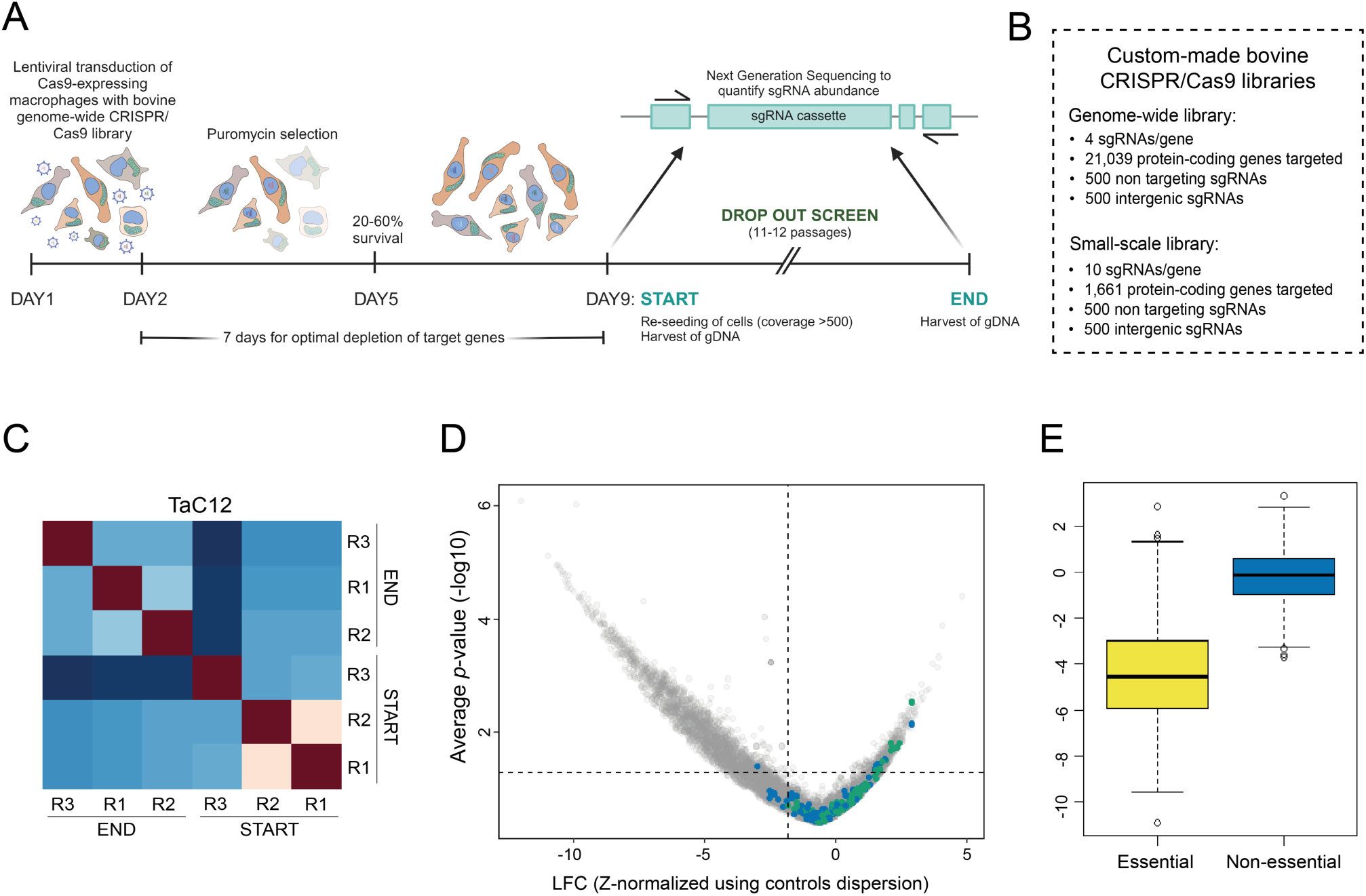
Genome-wide CRISPR/Cas9 screen in *Theileria annulata*-infected bovine macrophages identify host factors essential for parasite survival. **(A)** Schematic workflow of CRISPR dropout screens. Cas9-expressing infected macrophages (TaC12) were transduced with a pooled CRISPR sgRNA lentiviral library and maintained in culture for at least 11 passages. Genomic DNA was harvested at the START and END points and sgRNA abundance was quantified. The screen was performed in 3 biological replicates. **(B)** Detailed information on the bovine CRISPR/Cas9 libraries used in this study. **(C)** Correlation plot screen replicate. Dark red indicates high similarity between replicates, while dark blue indicates low correlation. **(D)** Volcano plot showing the hypergeometric distribution analysis of the TaC12 <u>genome-wide</u> screen. Each gene is plotted based on its average log_2_ fold change (LFC, z-normalized using control dispersion) and the negative log_10_ of its *p*-value. Intergenic and non-targeting controls are shown in blue and green, respectively. The horizontal dotted line corresponds to *p*-value = 0.05, and the vertical dotted line corresponds to average z-normalized LFC of the controls (intergenic and non-targeting) minus 1x standard deviation of the whole genetic screen. **(E)** Boxplot representing TaC12 genome-wide screen scoring (z-normalized LFC values) of groups of annotated “essential” and “non-essential” genes.

Infected macrophages were transduced with a Cas9 lentiviral construct. Protein expression was validated by Western blotting (**Figure S4A**). We also used a fluorescence based Cas9 reporter assay to select for cells expressing active Cas9 (**Figure S4B-C**). The genome-wide screen was performed at a coverage of 500 with three biological replicates (**Figure 3C**, **S4D-E**). Cells were transduced with lentiviruses for 24 h (DAY 1 -DAY 2, **Figure 3A**) aiming to achieve approximately 30% transduction efficiency, and positive cells were then selected with puromycin. We allowed the cells to grow for another four days to achieve optimal depletion of target genes. On DAY 9, one part of the puromycin-resistant cells was harvested (START point) and the other part reseeded to maintain coverage for later DNA extraction. The dropout screen continued for at least 11 passages (4 weeks), with cells kept under selection and reseeded after each passage. At the end of the screen, surviving cells were harvested (END point) and Illumina Next Generation Sequencing was performed to quantify the abundance of sgRNAs at END and START points. Log_2_ fold change (LFC) values were generated based on how sgRNAs changed in abundance throughout the screen, and the average LFC of three biological replicates was used for statistical analysis to identify bovine genes that were depleted in the dropout screen. The data were analyzed using a hypergeometric distribution, where gene LFC values were normalized using intergenic and non-targeting controls (**Table S2**) and displayed in a volcano plot (**Figure 3D**). A hit with a LFC < 0 indicates a reduction in cell fitness upon gene inactivation and consequently these cells are less abundant at the end of the screen compared to the beginning. We assessed the functionality of our library using a reference set of essential and non-essential genes identified by Hart and colleagues^36^ (**Figure 3E**). The reference genes were highly fitness conferring, while knockout of non-essential genes did not negatively affect TaC12 cell fitness. We then selected a list of ∼1,500 genes that significantly reduced cell fitness upon Cas9-mediated inactivation and designed a CRISPR sgRNA sublibrary (increasing the number of sgRNAs from 4 to 10 per gene) (**Figure 3B**) and performed a small-scale screen.

In this round of validation, we also screened the uninfected bovine macrophage cell line BoMac to identify genes essential for a cell type with a similar genetic background. Small-scale dropout screens were performed in biological triplicates following the protocol of the genome-wide CRISPR screen (**Figure 3A**, **S4G, I, Table S3**). Principal component analysis (PCA) of log-normalized counts confirmed the similarity of DAY 0 samples for both TaC12 and BoMac cell lines, as well as their resemblance to the plasmid DNA library (**Figure 4A**). The END samples of both cell lines were clearly separated from each other and from the DAY 0 samples. This suggests a divergent behavior of these two cell lines upon knockout out. While TaC12 screening results have a strong correlation among biological replicates, BoMac screens showed a lower replicate reproducibility, indicating a less consistent response in repeated experiments (**Figure 4B**). About 80% of candidate genes were validated as crucial for TaC12 survival (**Figure S4F**). However, 56% of the TaC12 *essentialome* is shared with BoMac (**Figure S4H**), suggesting that many of these genes are also involved in core biological processes. By removing the common fitness-conferring genes, we defined a list of 653 host genes that we considered to be the exclusive *Theileria*-infected macrophage *essentialome*, as these hits are dispensable for BoMac survival (**Figure 4C**, **Table S4**). One hundred genes that scored as non-essential in the genome-wide screen were included in the sublibrary and these grouped together with intergenic and non-targeting controls in both cell lines, highlighting the ability of our CRISPR library to consistently discriminate between essential and non-essential genes (**Figure 4C**, **Figure S4 F, H**).

**Figure 4.**
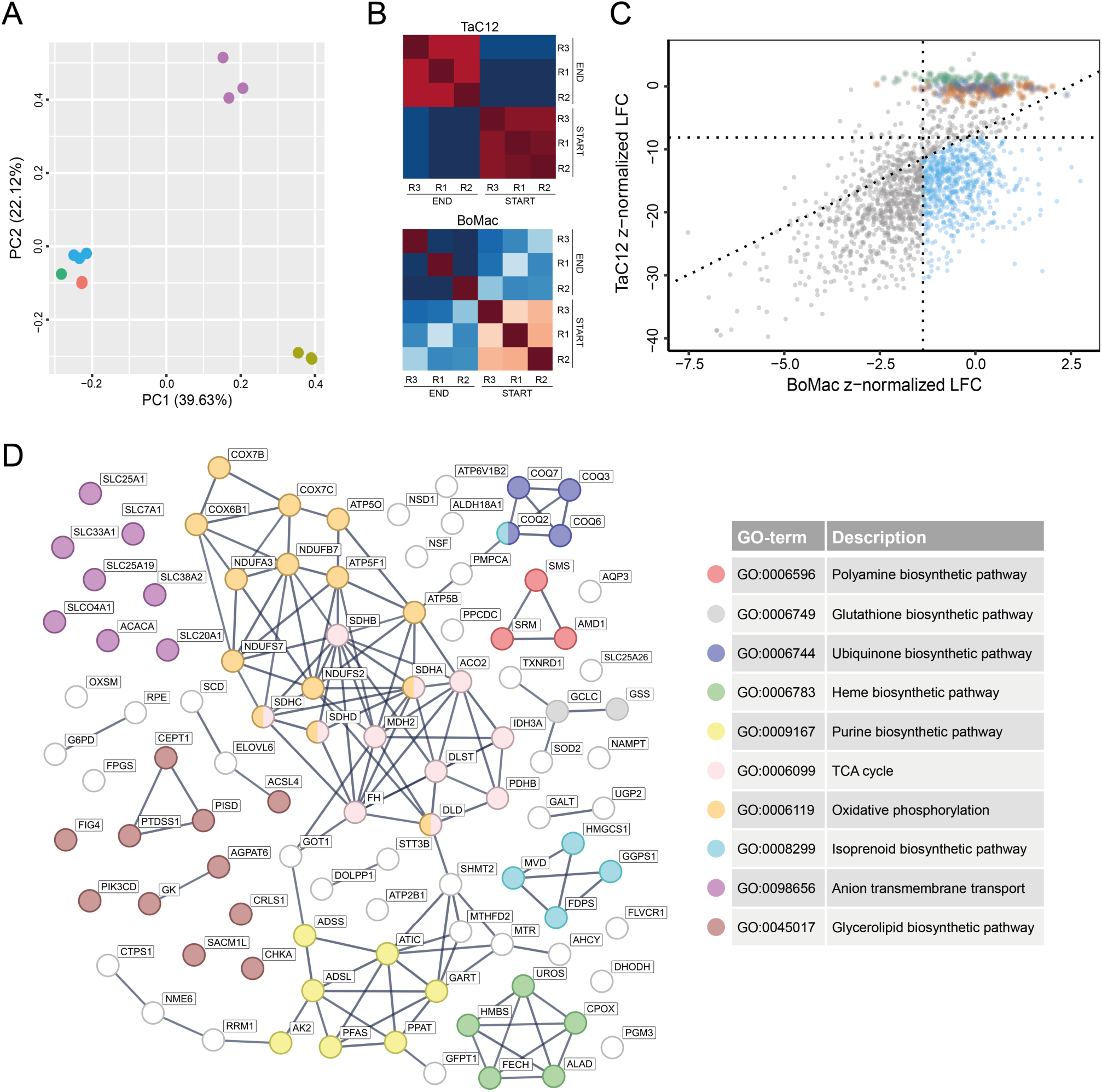
Pathway analysis of fitness-conferring host genes for *Theileria* schizonts. **(A)** Principal component analysis plot of the log-norm counts distribution of each biological replicate of small-scale screens in BoMac and TaC12. Color legend: BoMac_DAY0_r1-2-3 (blue), BoMac_END_r1-2-3 (purple), TaC12_DAY0_r1-2-3 (red), TaC12_END_r1-2-3 (sage green), CRISPR sublibrary (neon green). **(B)** Correlation plots of small-scale screen replicates. Dark red indicates high similarity between replicates, while dark blue indicates low correlation. **(C)** Scatter plot showing the results of the small-scale screen. Each gene is plotted based on z-normalized LFC values in infected versus uninfected cells. Genes depleted only in TaC12 cells are colored in light blue. Intergenic and non-targeting controls are shown in orange and green, respectively. One hundred non-essential genes from the TaC12 genome-wide screen are highlighted in orange. Dotted lines correspond to the average z-normalized LFC of the controls (intergenic and non-targeting) minus 1x standard deviation of each genetic screen. **(D)** Protein interaction network and Gene Ontology (GO) analysis of metabolic genes present in the *Theileria essentialome*. The list of 99 metabolic genes was analyzed using the STRING database (https://string-db.org/) on 19 July 2023. Line thickness indicates the strength of data support. Active interaction sources considered: text mining, experiments, databases, co-expression. Only interactions with the highest confidence score are shown. Gene ontology enrichment analysis identified 10 major host metabolic pathways.

### Host genes involved in proliferation, cell growth and metabolism are fitness-conferring in *Theileria*-infected bovine macrophages

To scrutinize the *Theileria essentialome* consisting of 653 bovine genes (**Table S4**) we performed a Gene Ontology analysis. This revealed numerous biological processes (**Table S5**) with a significant enrichment of proteins involved in ribosome biogenesis and assembly (GO:0042254), mRNA metabolism (GO:0016071), RNA processing (GO:0006396), translation (GO:0006412) and cell cycle progression (GO:0022402), highlighting the importance of continuous protein synthesis to cope with the proliferative phenotype induced by *T. annulata*. Genes involved in the host cell metabolism make up 15% of the *Theileria* essentialome, as revealed by STRING analysis (**Figure 4D**), including biosynthesis of isoprenoids (GO:0008299), glycerolipids (GO:0045017), polyamines (GO:0006596), glutathione (GO:0006749), heme (GO:0006783) and ubiquinone (GO:0006744). The tricarboxylic acid cycle is the most enriched metabolic process in our dataset according to KEGG pathway analysis (**Figure S5A**). For example, mitochondrial enzymes such as aconitase (*ACO2*), succinate dehydrogenase (SDH subunits A-D), fumarate hydratase (*FH*), malate dehydrogenase (*MDH2*) and the catalytic subunit of isocitrate dehydrogenase 3 (*IDH3A*) are exclusively fitness-conferring for TaC12 (**Figure S5B**). As expected, since the entire phylum of apicomplexan parasites lacks the *de novo* purine biosynthesis pathway, the *Theileria essentialome* contains several genes involved in the biosynthesis of purine precursors (GO:0009167) (**Figures 4D**, **S6A**). Most of these genes catalyze consecutive reactions in the generation of inosine monophosphate, the intracellular precursor of adenosine and guanosine monophosphates (AMP, GMP). Interestingly, while enzymes that convert IMP to AMP are present (*ADSS* and *ADSL*), those deputed to synthesize GMP (*IMPDH* and *GMPS*) are absent, highlighting a specific requirement for adenine nucleotides by infected cells (**Figure S6B**).

### Host biosynthesis of glutathione, polyamines, and heme is critical for *Theileria annulata*-infected leukocytes

Among the metabolic hits from the *Theileria* screen we selected three pathways for validation by reverse genetics and chemical inhibition. We aimed for metabolic processes that are dispensable for the host and affect *Theileria*-infected cells through interference with the parasite nutrients supply. Within the glutathione biosynthetic pathway, glutathione is a ubiquitous intracellular and extracellular antioxidant tripeptide synthesized by a two-step reaction catalyzed by the heterodimeric enzyme γ-glutamylcysteine synthetase (γ-GCS; *GCLC*, catalytic subunit; *GCLM*, regulatory subunit) and glutathione synthetase (GSS) (**Figure 5A**). In both genome-wide and small-scale screens, perturbation of the *GSS* and *GCLC* significantly reduced the survival of infected cells over time, while uninfected macrophages showed no defect in cell survival (**Figure 5A**). A single CRISPR/Cas9 knockout of bovine *GSS* (**Figure 5A**, **S7A**), resulted in a population of BoMac with 58.7% frameshift mutations in the target gene locus, while >90% of TaC12 retained the wild-type *GSS* sequence. This result suggests that TaC12 *GSS* mutants are not viable and undergo cell death upon gene knockout. To support these data, we treated cells for 72 h with different concentrations of buthionine sulfoximine (BSO), a well-established irreversible inhibitor of γ-GCS,^37^ and observed that TaC12 were more sensitive than BoMac in the 30 - 100 µM BSO range (**Figures 5A**, **S7C**), confirming the importance of glutathione biosynthesis exclusively for infected cells. Polyamines are molecules derived from amino acids that are important for cell proliferation and protein synthesis. They are important for rapidly proliferating cells. Mammalian cells have a classical forward-directed *polyamine biosynthetic pathway* that produces polyamines from arginine and methionine, but they also can convert spermine back to spermidine and putrescine via a retroconversion pathway (**Figure 5B**). One of the top depleted genes in the TaC12 screen was spermidine synthase (*SRM*), which was dispensable for BoMac (**Figure 5B**). Targeted *SRM* CRISPR mediated knockout resulted in >82% frameshift mutations in BoMacs, while no significant mutations were detected in surviving TaC12 cells (**Figures 5B**, **S7A**). Consistent with this finding we observed that when cells were treated with the nitric oxide donor sodium nitroprusside (SNP), a potent inhibitor of polyamine biosynthesis,^38^ TaC12 showed a >70% reduction in cell viability compared to BoMac (**Figures 5B**, **S7D**). Ablation of five out of the eight enzymes (**Figure 5C**) of the host *heme biosynthetic pathway* negatively affected the fitness of TaC12 cells in the screen, while no phenotype was observed in uninfected macrophages (**Figure 5C**). We selected *HMBS* for further validation. No viable knockout could be achieved in *Theileria*-infected cells, whereas we were able to generate a polyclonal population of BoMac cells that acquired frameshift mutations in the target gene (**Figure 5C**, **S7A**). Treatment with salicylic acid (SA), an inhibitor of FECH,^39^ reduced the viability of TaC12 cells by >60%, whereas BoMac viability remained unchanged (**Figure 5C**, **S7E**). Together, these data validate the critical role of the host glutathione, polyamine, and heme biosynthesis in *Theileria*-infected cells.

**Figure 5.**
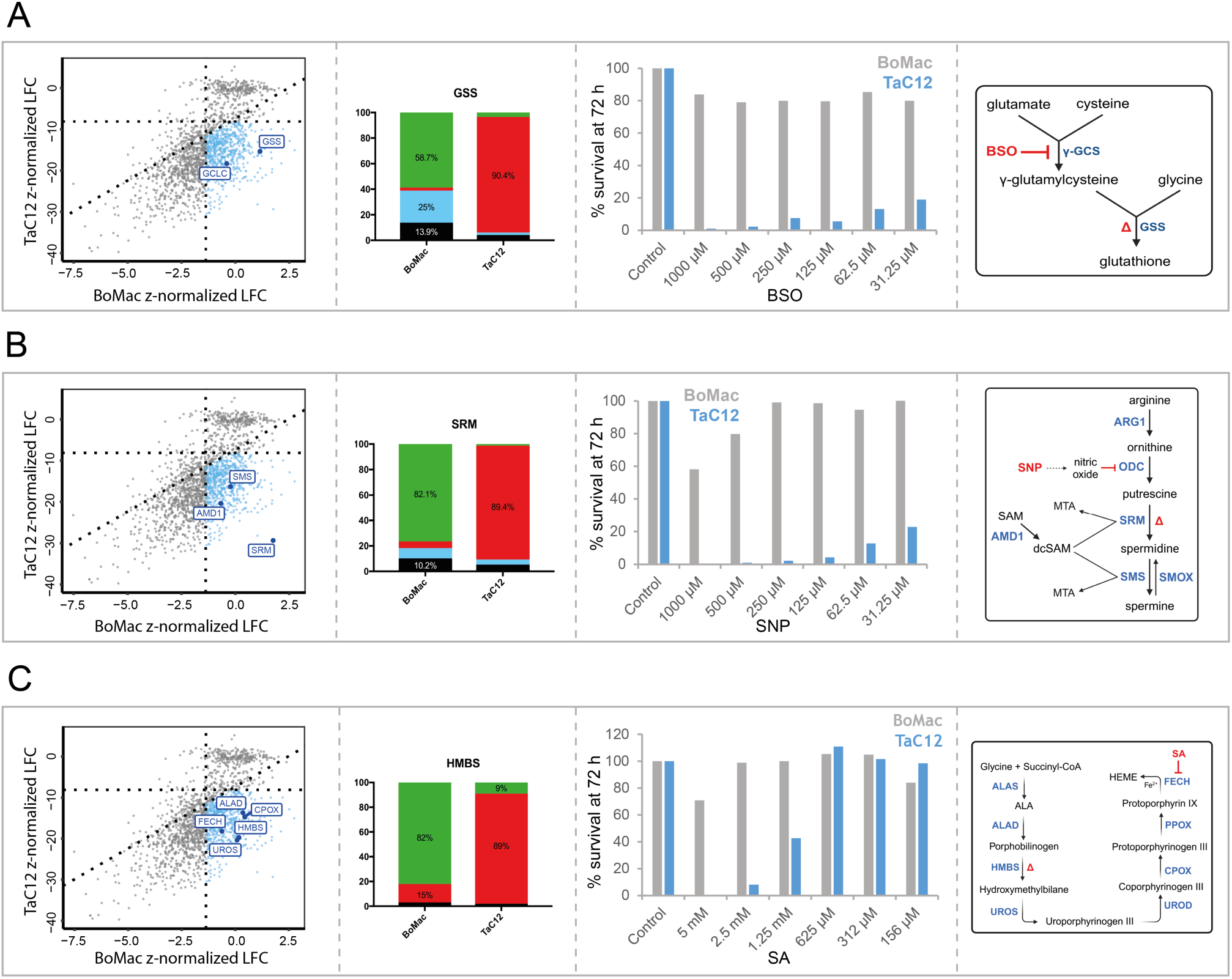
Host glutathione, polyamine, and heme pathways are essential for *Theileria*. **A)** Scoring of *GCLC* and *GSS* in TaC12 vs. BoMac screens. TIDE analysis of *GSS* CRISPR/Cas9 knockout in BoMac and TaC12 cells. Resazurin viability assay showing the percentage of survival of BoMac and TaC12 cells after 72 h treatment with buthionine sulfoximine (BSO), an inhibitor of γ-GCS, as indicated in the schematic of the glutathione biosynthetic pathway. The gene knocked out in TaC12 and BoMac is indicated with delta (Δ) in the pathway. **(B)** Scoring of *AMD1*, *SMS* and *SRM* in TaC12 vs BoMac screens. TIDE analysis of *SRM* CRISPR/Cas9 knockout in BoMac and TaC12 cells. Resazurin viability assay showing the percentage of survival of BoMac and TaC12 cells after 72 h treatment with sodium nitroprusside (SNP), an inhibitor of ODC as indicated in the schematic of the polyamine biosynthetic pathway. The gene knocked out in TaC12 and BoMac is indicated with delta (Δ) in the pathway. **(C)** Scoring of *ALAD*, *HMBS*, *UROS*, *CPOX* and *FECH* in TaC12 vs BoMac screens. TIDE analysis of *HMBS* CRISPR/Cas9 knockout in BoMac and TaC12 cells. Resazurin viability assay showing the percentage of survival of BoMac and TaC12 cells after 72 h treatment with salicylic acid (SA), an inhibitor of FECH as indicated in the schematic of the heme biosynthetic pathway. TIDE and viability experiments were performed in triplicate with similar results (Fig. S4 E-G-H-I). A representative replicate is shown for each. The gene knocked out in TaC12 and BoMac is indicated with delta (Δ) in the pathway.

### *Plasmodium* and *Theileria* schizont development depend on several shared host pathways

*Plasmodium* liver-stage infection remains one of the least characterized but most attractive targets for malaria prevention as it represents a bottleneck in parasite development with only very few parasites arriving and successfully establishing in hepatocytes. Experimental limitations, such as the low infection rate of *Plasmodium,* prompted us to seek an alternative approach to study the host interplay of the hepatic schizont stages of the human malaria parasite and to search for common essential host metabolic pathways of hematozoans. To this end, we focused on the 99 identified metabolic enzymes from the *Theileria essentialome* of *Theileria*-infected cells (**Figure 4D**) and searched for overlaps with *P. falciparum* hepatocyte gene essentiality predictions in a rich and compromised extra-parasitic environment of the host cell, respectively (**Figures 2E-F**, **6**). We identified 24 enzymes, including genes encoding mitochondrial complex II subunits (*SDHA*, *SDHB*, *SDHC*, *SDHD*) and genes involved in polyamine (*AMD1*, *SRM*) and glutathione biosynthesis (*GCLC*, *GSS*), as essential for both parasites when *Plasmodium* grows in a rich environment (**Figure 6**). Of these only seven genes, belonging to the heme and purine biosynthetic pathways, were identified as essential for both parasites when *Plasmodium* growth was modelled in a compromised environment (**Figure 2F**). This suggests that when metabolites such as polyamines and glutathione are available for uptake in the host cytosol, *Plasmodium* parasites scavenge these nutrients. However, in the absence of these molecules in a compromised environment, our model suggests that parasite *de novo* biosynthesis is enough to produce sufficient metabolites. Malaria parasites encode all the genes necessary to synthesize both polyamines and glutathione, but would probably save energy by scavenging from the host. On the other hand, all apicomplexans have lost the ability to synthesize purine nucleotides endogenously and rely on the mammalian host as a source of purines^40^. As expected, we found that the host genes of *ATIC*, *GART*, *PFAS* and *PPAT* are highly fitness-conferring for *Theileria* and essential for all *P. falciparum parasitosomes* (**Figure 6**). We also identified the heme biosynthesis pathway, represented by the host genes *CPOX*, *HMBS* and *UROS*, as essential for both parasites under all conditions. Essentiality of the host heme pathway for *Theileria*-transformed macrophages was not surprising since *Theileria* lacks all enzymes of the heme pathway.^41^ However, the prediction that host HMBS is essential for the *P. falciparum* was unexpected, since *Plasmodium* possesses the enzymatic machinery for heme biosynthesis.^41^

**Figure 6.**
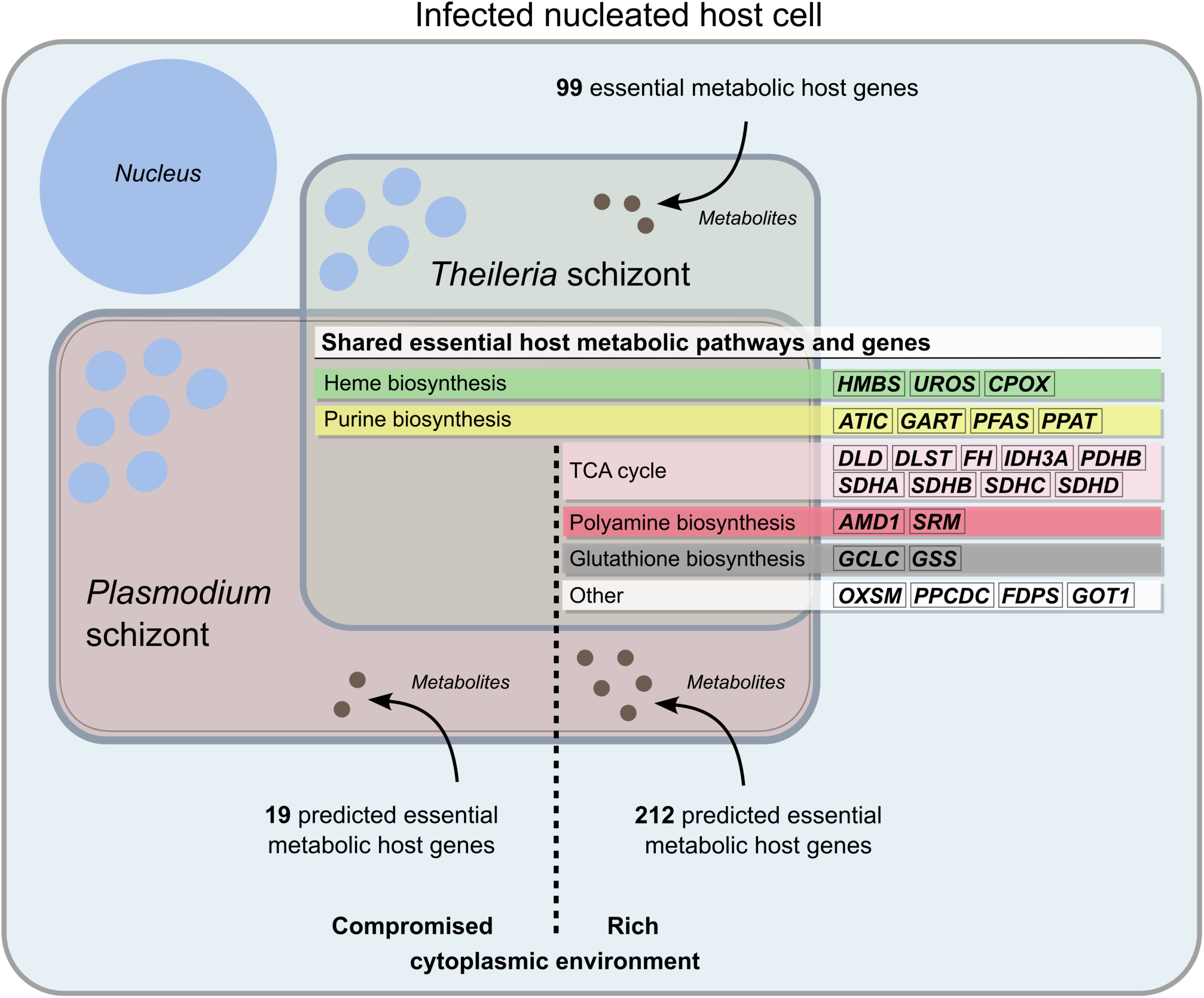
Host metabolic pathways and genes essential for *Plasmodium* and *Theileria* schizont survival in nucleated host cells. Venn diagram showing multinucleated parasite schizonts and the overlap between the *Theileria* metabolic essentialome and *P. falciparum* host metabolic gene essentiality predictions in a compromised and rich extra-parasitic environment.

### Perturbation of environmental heme pools strongly influences *Plasmodium* schizont development

The importance of the host hepatic heme biosynthesis pathway has not been thoroughly investigated for *P. falciparum*, possibly because malaria parasites are considered heme prototrophic eukaryotes.^41^ Transcriptomic data of *P. falciparum* liver stages are not available but have been thoroughly studied for the rodent model parasite of *P. berghei*. *P. berghei* heme biosynthesis gene expression data indicate low activity of the endogenous pathway during biomass increase in the liver stage^42^ (**Figure 7A**). Interestingly, *Pb*FECH, the enzyme that catalyzes the final step of the pathway, is significantly upregulated, with expression peaking at the late exo-erythrocytic stage (54 & 60 hpi). Previous research has shown that *Pb*FECH knockout parasites grow to only half the size compared to wild-type parasites.^43^ This suggests that parasites may use, at least in part, host cell-derived or systemically available porphyrins. Notably, our models suggest that the mammalian heme pathway plays a central role in the development of the *Plasmodium* exoerythrocytic stage, with three heme pathway genes being identified as potentially essential for both *Plasmodium* and *Theileria*. We chose to knock out *HMBS*, the most upstream gene, to inactivate the entire pathway. For the knockout experiment, we used the haploid cell line HAP1 because knockout approaches are very efficient in these cells, and for infection, we used the rodent model parasite *P. berghei* because infectious sporozoites are readily accessible and able to infect HAP1 cells. Successful ablation of *HMBS* and complementation in HAP1 cells was confirmed at both the gene and protein level (**Figures 7B**, **S7F**). Although HAP1 ΔHMBS cells are unable to synthesize heme *de novo*, they may be able to import extracellular porphyrins. Fetal calf serum (FCS) added to mammalian cell culture medium contains residual concentrations of heme (∼1 -2µM).^44^ We therefore depleted heme and could show that clonal *HMBS*-deficient HAP1 cells do not grow in heme-depleted media, and that this growth defect can be restored by hemin supplementation (**Figure 7C**). We then investigated whether *P. berghei* parasites can successfully develop in HAP1 cells with impaired heme metabolism. We infected six individual HAP1 ΔHMBS clones with *P. berghei* sporozoites and compared parasite size at 48 hpi. As predicted by our metabolic model, we observed a defect in intracellular parasite growth, as parasites were significantly smaller compared to those in WT cells (**Figure 7D**). The marked reduction in parasite size was common to all infected ΔHMBS clones. Next, we analyzed parasite growth after *HMBS* complementation. The reduced size phenotype observed in the parental knockout cells was rescued (**Figure 7E**), indicating that reactivation of the interrupted host heme biosynthetic pathway allows the successful growth of *Plasmodium* parasites.

**Figure 7.**
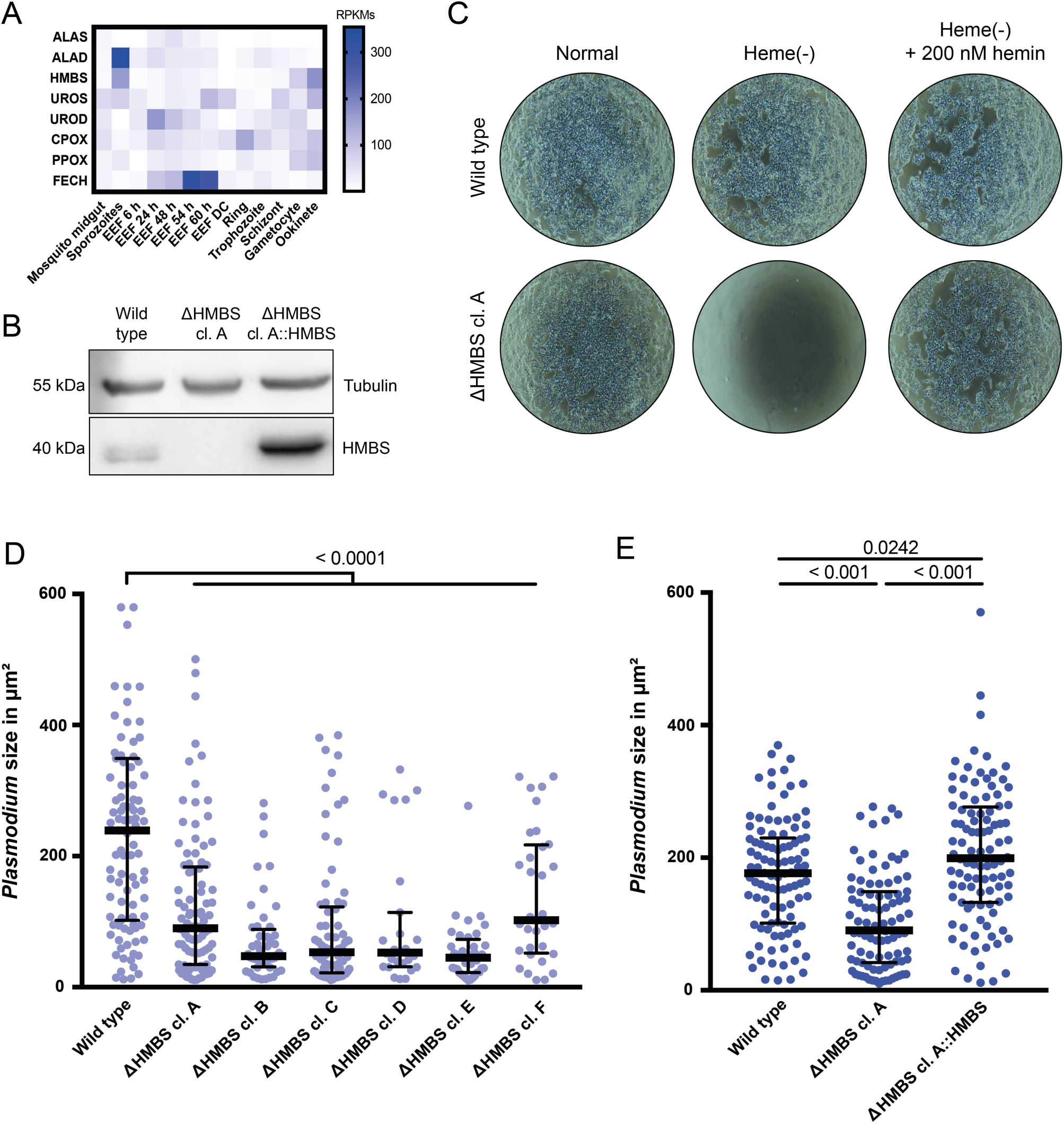
Environmental heme from the host cell plays a key role in the development of *Plasmodium* exo-erythrocytic schizonts. **(A)** Expression of *P. berghei* heme biosynthesis genes throughout the parasite’s life cycle. Most parasite genes are expressed at low levels during the liver stage (EEF), except for *FECH*. EEF, exo-erythrocytic form; DC, detached mammalian cell containing merosomes. (**B)** Western blot of whole cell lysates from HAP1 wild type, ΔHMBS clone (cl.) A, and a pool of ΔHMBS cl. A::HMBS complemented cells. Tubulin was used as a loading control. **(C)** Phase contrast images of HAP1 wild type and HAP1 ΔHMBS cl. A after 48 h incubation in normal 10% FCS-IMD media, heme-depleted 10% FCS-IMDM, and the latter with hemin supplementation. (D) *P. berghei* exo-erythrocytic schizont size at 48 h pi in HAP1 WT and six HAP1 ΔHMBS clones A - F. **(E)** Parasite size at 48 hpi in HAP1 WT, ΔHMBS cl. A, and ΔHMBS cl. A::HMBS complementation. *P*-values are shown in D and E.

## DISCUSSION

Apicomplexan parasites, due to their inherent metabolic constraints, rely heavily on scavenging essential metabolites from their hosts.^40,41,45^ This intricate metabolic interplay between *Plasmodium* or *Theileria* becomes particularly critical prior to erythrocyte invasion, when the parasites expand extensively within their nucleated host cells. By employing a synergistic approach that combines genome-scale metabolic network modeling of *P. falciparum* infected hepatocytes and genome-wide CRISPR/Cas9 screening of *Theileria-* infected macrophages, we have gained critical insights into the shared metabolic dependencies during the early life-cycle stages within the mammalian host, and pinpointed the metabolic genes of the infected host cell that are critical for parasite survival.

Because the clonal proliferation of *Theileria*-transformed cells makes them ideal for large-scale screens that are difficult to perform in *Plasmodium*-infected hepatocytes, we performed a CRISPR dropout screen in *Theileria*. We identified a set of 653 host genes, referred to as the *Theileria essentialome*, which significantly impact the growth of *Theileria*-infected cells. This resource includes not only genes involved in protein synthesis and cell proliferation, but also comprises a significant enrichment of host metabolic genes. Since the survival of *Theileria*-transformed macrophages depends on the presence of a viable parasite,^46^ we infer that much of the *Theileria essentialome*, particularly those genes involved in metabolic processes, is required for parasite survival. Twenty-four host metabolic enzymes of the *Theileria essentialome* (e.g., of the TCA cycle, polyamine, glutathione, heme, and purine biosynthesis) are shared with the predicted *P. falciparum* hepatocyte model (summarized in Figure 6). Of these, seven genes belonging to the heme biosynthesis and purine synthesis pathways were predicted to be essential for *Plasmodium* growth under all modeled metabolic conditions. This highlights the need to consider the parasite’s environment, including the host cell cytoplasm and the extracellular milieu surrounding the host cells, when assessing host-parasite metabolic dependencies. For example, our predictions show that *Plasmodium* schizonts preferentially rely on host glutathione and polyamine synthesis in a “rich” nutrient environment but can switch to endogenous synthesis when the availability of metabolites in the host is limited.

An important metabolite in the glutathione pathway is GSH, which plays a critical role in protecting cells from oxidative stress. *Plasmodium* has an active GSH biosynthetic pathway that is vital for its development in the mosquito. Since *P. berghei* GSS knockout mutants develop efficiently in the liver *in vivo*,^16^ this supports our prediction of a preferential dependency on host *GSS*. While *Plasmodium* can potentially rely on its own glutathione synthesis pathway when needed, our data indicate that modulating host GSH levels, in combination with antimalarial therapy, may be a promising line of future research in malaria therapy. Interestingly, knockdown of host *GSS* has previously been reported to increase the susceptibility of human macrophages to the anti-leishmanial activity of glucantime.^47^ In addition, targeting GSH in the context of anti-parasite therapy has parallels to anti-cancer therapy, where high GSH levels are also essential for cancer cell survival. Strategies such as inhibiting GSH synthesis by cysteine starvation or targeting of the amino acid antiporter system Xc-and combination treatment with the γ-GCS inhibitor buthionine sulfoximine (BSO) have already shown promise as anti-tumor combination therapies, ^48–50^ and warrant exploration in the context of malaria and theileriosis therapies.

Another important finding is the essentiality of host *SRM* for the *Plasmodium* liver stage development, contingent on the environment. Polyamines are essential molecules for all eukaryotes, and *Plasmodium* parasites are unique among apicomplexans in encoding a complete set of enzymes for endogenous polyamine biosynthesis. *Theileria*, on the other hand, is completely auxotrophic for these amines.^51–53^ Malaria parasites can synthesize all three major polyamines, including spermine, although their genome lacks an orthologue of the SMS gene.^53^ Instead, they encode for a promiscuous SRM enzyme capable of producing spermidine and, to a much lesser extent, spermine.^54^ The physiological role of spermine for *Plasmodium* parasites remains unclear, and there is no evidence for an uptake mechanism, also warranting further investigation.

Our data also reveal the shared dependency of both *Plasmodium* and *Theileria* on four host genes (*ATIC*, *GART*, *PFAS* and *PPAT*) involved in purine synthesis. This observation is consistent with the fact that the entire phylum of Apicomplexan parasites lacks an endogenous purine synthesis pathway, highlighting their dependency on the acquisition of these bases from the mammalian host.^40^

In contrast, diverse heme acquisition strategies have been described among Apicomplexan parasites. While *Plasmodium* and *Toxoplasma* possess a complete and functional *de novo* heme biosynthetic pathway, parasites such as *Babesia*, *Theileria* and *Cryptosporidium* depend on acquiring heme from the host cell.^41^ Our experiments provided insights into the consequences of loss of the host heme pathway in *P. berghei* and confirmed the unexpected predictions of our model. When *HMBS* was compromised, *P. berghei* failed to reach its normal size at 48 hpi. Strikingly, complementation of *HMBS* completely reversed this phenotype. While there are conflicting data regarding the essentiality of *Pb*ALAS in the liver, external ALA could rescue *P. berghei* lacking *ALAS*.^55,56^ This is consistent with the idea that liver-stage *Plasmodium* can utilize external ALA for heme synthesis. Rathnapala et al. further demonstrated that FECH-deficient *P. berghei* grew smaller in size during liver stage development.^43^ Taken together, this suggests that endogenous heme synthesis becomes critical during liver development, in contrast to the intraerythrocytic stages.^55–58^ Our results now add to this picture by suggesting that the host heme synthesis enzymes upstream of FECH play a fitness-conferring role in the parasite development. However, the identity of porphyrin transporters in apicomplexans remains relatively unexplored,^41^ and requires further investigation.

In conclusion, our study has demonstrated the power of integrating computational modeling with experimental approaches to uncover crucial host genes essential for intracellular survival of two of the deadliest parasites on the planet. The methods developed in this study provide an adaptable framework for investigating a range of other host-parasite interactions, allowing a deeper understanding of common host dependencies. Moreover, our findings open up exciting opportunities to simultaneously target both host and parasite pathways and to explore gene essentiality in specific environmental contexts and cellular niches, paving the way for innovative combination therapies against these devastating pathogens.

## ACKNOWLEDGEMENTS

We thank the Department for BioMedical Research (DBMR) and the Flow Cytometry and Cell Sorting Facility (FCCS) of the University of Bern, Switzerland. Financial support came from the Swiss National Science Foundation (CRSII5_198543, 310030_189127, PZ00P3_173972). Figures were created with the help of BioRender.

## AUTHOR CONTRIBUTIONS

Conceptualization, KW, SR, PO; methodology, JGD, KW, VHr, VHa, SR, PO; investigation, MMr, MMs, KW, RC, AN, DJ, JZ, MB; formal analysis, MMr, MMs, KW, RC, AN, MGF, PO; writing – original draft, MMr, MMs; writing – review & editing, KW, RC, JGD, AN, VHr, VHa, SR, PO; funding acquisition, VHa, VHr, SR, PO; resources, VHa, VHr, SR, PO; supervision, VHa, VHr, SR, PO

## STAR ★ METHODS

### KEY RESOURCES TABLE<colcnt=3>

**KEY RESOURCES TABLE**
| REAGENT or RESOURCE | SOURCE | IDENTIFIER |
| --- | --- | --- |
| **Deposited Data** |  |  |
| Human GEM Recon 3 | Brunk et al, 2018 | https://www.vmh.life |
| Thermodynamically curated GEM Recon 3D | Masid et al. 2020 | https://github.com/EPFL-LCSB/redhuman |
| Human protein data | The Human Protein Atlas | https://www.proteinatlas.org/ |
| Fluxomics data for hepatocyte model | Agren et al. 2014 | doi:10.1002/msb.145122 |
| Metabolomics data for hepatocyte model | Gille, Bolling, Hoppe, et al. 2010 | doi:10.1038/msb.2010.62 |
| **Experimental Models: Cell Lines** |  |  |
| TaC12 | Shiels et al., 1992 | doi:10.1242/jcs.101.1.99 |
| BoMac | Stabel et al., 1995 | doi:10.1016/0165-2427(94)05348-v |
| HAP1 | Carette et al., 2011 | doi:10.1038/nbt.1857 |
| **Software and Algorithms** |  |  |
| MATLAB | MathWorks | R2021b |
| CPLEX | IBM | CPLEX Optimization Studio V12.10.0 |
| matTFA | Salvy, et al. 2018 | https://github.com/EPFL-LCSB/mattfa |
| lumpGEM | Ataman and Hatzimanikatis, 2017 | https://github.com/EPFL-LCSB/redgem |
| **Other** |  |  |
| Repository with code for model reconstruction and metabolic analysis | This paper | https://github.com/EPFL-LCSB/host_parasite_interactions |

## RESOURCE AVAILABILITY

### Lead contact

Further information and requests for resources and reagents should be directed to the lead contact, Philipp Olias.

### Materials availability

All unique/stable reagents generated in this study are available from the Lead Contact with a completed Materials Transfer Agreement.

### Data and code availability

The code generated during this study is available at https://github.com/EPFL-LCSB/host_parasite_interactions.

## METHOD DETAILS

### Hepatocyte specific model

Starting with the thermodynamically curated human genome-scale Recon 3D^31,32^, we reconstructed a hepatocyte metabolic model by taking into account the physiology of hepatocytes and the genes expressed in liver cells. Towards this end, we defined the physiology of hepatocytes by integrating in the human Recon3D model publicly available fluxomics data for 92 boundary reactions^29^ and metabolomics data for 213 metabolites^30^ from previous hepatocyte model reconstructions. Additionally, we set a growth rate of the hepatocyte to a maximum of 0.014 h^-1^ corresponding to a doubling time of 49.5 h^59^ and the ATP maintenance rate to at least 1.07 mmol/gDW/h^60^. The Human Protein Atlas (www.proteinatlas.org) ^28^ was used to identify 1853 metabolic genes present in liver cells. By mapping these genes to the Recon 3D reactions (using the gene-protein-reaction rules), 5926 reactions conformed the starting core of the hepatocyte model. We then formulated the following MILP to integrate additional reactions required for the core reactions to carry flux:

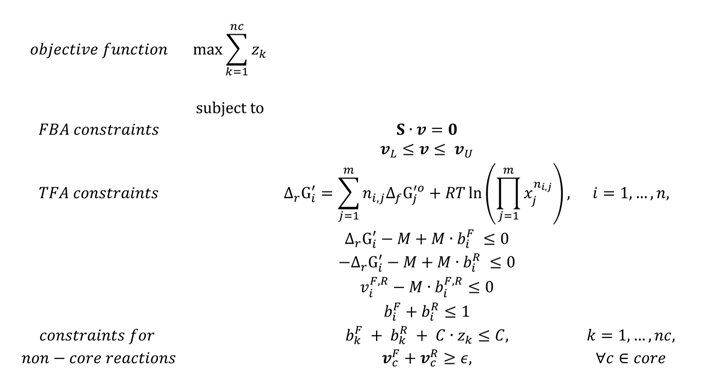

where *m* is the number of metabolites in the network, *n* is the number of reactions in the network, *nc* is the number of non-core reactions, *z*_*k*_ are binary variables for all the non-core reactions, ***S*** is the stoichiometric matrix, ***v*** are the net fluxes for all the reactions and *v*^*F*^_*j*_, *v*^*R*^_*j*_ are the corresponding net-forward and net-reverse fluxes, so that, *v*_*i*_ = *v*^*F*^_*j*_ − *v*^*R*^_*j*_, *for all i* = 1, …, *n*. ***v***_%_ and ***v***_&_ are the lower and upper bound, respectively, for all the reactions in the network. Δ**G**_’_ (is the Gibb’s free energy of the reactions and **b**^*F*^ and **b**^*R*^ are the binary variables for the forward or reverse fluxes of all the reactions (coupled to TFA). *M* is a big constant (bigger than all upper bounds), *C* is an arbitrary large number, and *ε* a small number. With this formulation, if *z*_*k*_ = 0, then reaction *k* is active.

As a result, the hepatocyte model contained a total of 3899 metabolites, 6209 reactions, and 1927 genes.

### Liver-specific model for *Plasmodium falciparum*

The liver-iPfa model is reconstructed from the published iPfa genome-scale model for *P. falciparum*^25^ by blocking the reactions that transport in and out of the parasite compounds that are not present in the liver model. By mapping the extra-parasitic metabolites from iPfa to the cytosolic metabolites from the hepatocyte model, we found that from the 241 nutrients that the parasite can transport, 165 (**Table S1**) are present in the cytosol of the liver model. We then blocked the corresponding 76 transport reactions and removed the reactions that could not carry flux in iPfa, resulting in a liver-iPfa model of 737 metabolites, 889 reactions, 210 genes, and a growth of 0.073 h-1 (doubling time of 9.48 h).

### Minimal nutritional requirements for liver-stage *Plasmodium falciparum*

In order to investigate the minimal set of nutrients that the liver-stage *P. falciparum* needs to sustain growth, we applied the *in silico minimal medium* (iMM) method^31^ to the liver-iPfa model. The iMM method identifies the minimal set of nutrients that are required to satisfy a specific growth rate. In this case, we identified that parasites need a minimum of 31 nutrients to grow. Given the flexibility of the intracellular metabolism to synthesize the required precursors for biomass, there exist alternative sets of nutrients that could serve to achieve the same growth rate. In this case, using iMM we found 1792 alternative sets of 31 nutrients, leading to a total of 47 unique nutrients across sets. By classifying the presence of these nutrients in the corresponding 1792 sets, we identified 23 nutrients that appear in all sets, which we call constitutive, and 24 nutrients that are not appearing systematically in all sets because they can substitute each other, as they contribute with the same backbone moieties.

### The *parasitosome* as a representation of the metabolic state of *Plasmodium falciparum*

The *parasitosomes* are generated using the lumpGEM method^61^, setting as objective the biomass reaction. This includes three steps (**Figure S1A**): ^62^ definition of the extra-parasitic environment in the metabolic model, (2) identification of the minimal number of reactions required for growth, the resulting set of reactions is called minimal network (MiN), and (3) generation of an individual reaction by collapsing all the reactions present in the MiN into one lumped reaction. As a result, the *parasitosome* reaction represents the interactions of the parasite with its environment, containing nutrients as substrates and secretions as products in the stochiometric equation. Given the flexibility of the intracellular metabolism, for the same physiological conditions, several MiNs can exist, representing alternative metabolic pathways that the parasites can use to synthesize the biomass precursors.

We have formulated *parasitosomes* for the two case studies with rich or compromised extra-parasitic environment (ePE). In the case that a rich ePE is available, a minimum of 248 reactions are required and there exist 151 alternative MiNs (**Figure S1C**). By lumping the MiNs into individual reactions, we generated 151 *parasitosomes* that use a total of 75 nutrients and secrete a total of 27 products (**Figure S1D**). In the case where the parasite encounters a compromised extra-parasitic environment, the size of the MiNs is 307 reactions and there exist 8 alternatives (**Figure S1C**), resulting in 8 alternative *parasitosomes* that take up a total of 36 nutrients and secrete 36 products (**Figure S1D**).

### Reconstruction of a hepatocyte-parasite integrated metabolic model

The generated *parasitosome* reactions were integrated into the hepatocyte metabolic model. We then evaluated whether the bounds of the transport reactions in the hepatocyte model needed to be modified in order to sustain the growth of both the parasite and the hepatocyte. To do this, we formulated the following MILP:

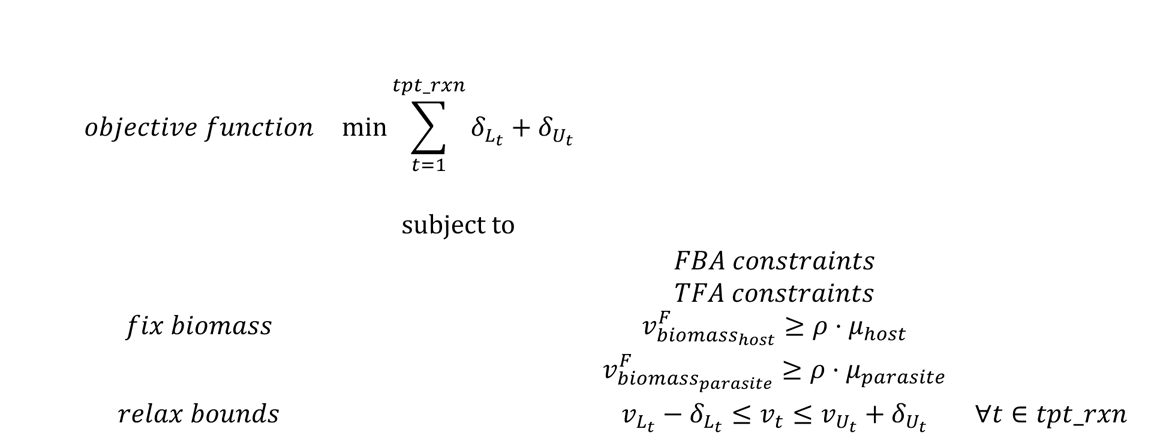

where *tpt_rxn* is the set of transport reactions in the network, ***v*** are the net fluxes for all the reactions and, ***v***_%_ and ***v***_&_ are the lower and upper bound, respectively, for all the reactions in the network. *δ*_*Lt*_ and *δ*_*Ut*_ are the relaxations of the lower and upper bound respectively. *μ*_*host*_ and *μ*_*parasite*_ are the theoretical maximum biomass for host and parasite, respectively and, *ρ* is a percentage (0 - 100%).

As a result, the integrated model contains 3906 metabolites, 6377 reactions, and 1927 genes.

### Gene essentiality analysis in the integrated hepatocyte-parasite model

We performed gene essentiality analysis^63^ in the healthy hepatocyte model and in the integrated hepatocyte-parasite model. In this analysis, hepatocyte genes are knocked out individually in the network. When a gene encoding for an enzyme is knocked out, the reaction rate is set to zero and we evaluated whether the corresponding KO influences biomass production.

For the integrated hepatocyte-parasite model, we knocked out the hepatocyte genes and we evaluated the impact on biomass production for the hepatocyte and for each individual *parasitosome*. We then selected the genes that were essential for parasite growth but not for hepatocyte biomass production, nor for the healthy hepatocyte.

### Cell culture

*T. annulata*-infected macrophages (TaC12) were maintained in L15 medium (Gibco, Thermo Fisher Scientific, Waltham MA, USA), while uninfected bovine macrophages (BoMac)^64^ and human embryonic kidney cells (HEK293FT) were grown in DMEM medium (Gibco). HAP1 cells, a human near-haploid cell line derived from chronic myelogenous leukemia KBM-7 cells, were maintained in IMDM medium (Gibco). All media were supplemented with 10% heat-inactivated fetal calf serum (BioConcept, Allschwill, Basel, Switzerland), 2 mM L-glutamine (BioConcept), 10 U/mL penicillin-streptomycin (BioConcept). L15 and DMEM media were also supplemented with 10 mM HEPES pH 7.2 (Merck). All cell lines were grown at 37 °C in an atmosphere containing 5% CO_2_, except for TaC12, which were cultured at 0% CO_2_.

### Plasmids

All plasmids used in this study are listed in the Table S6.

**Table S6.**
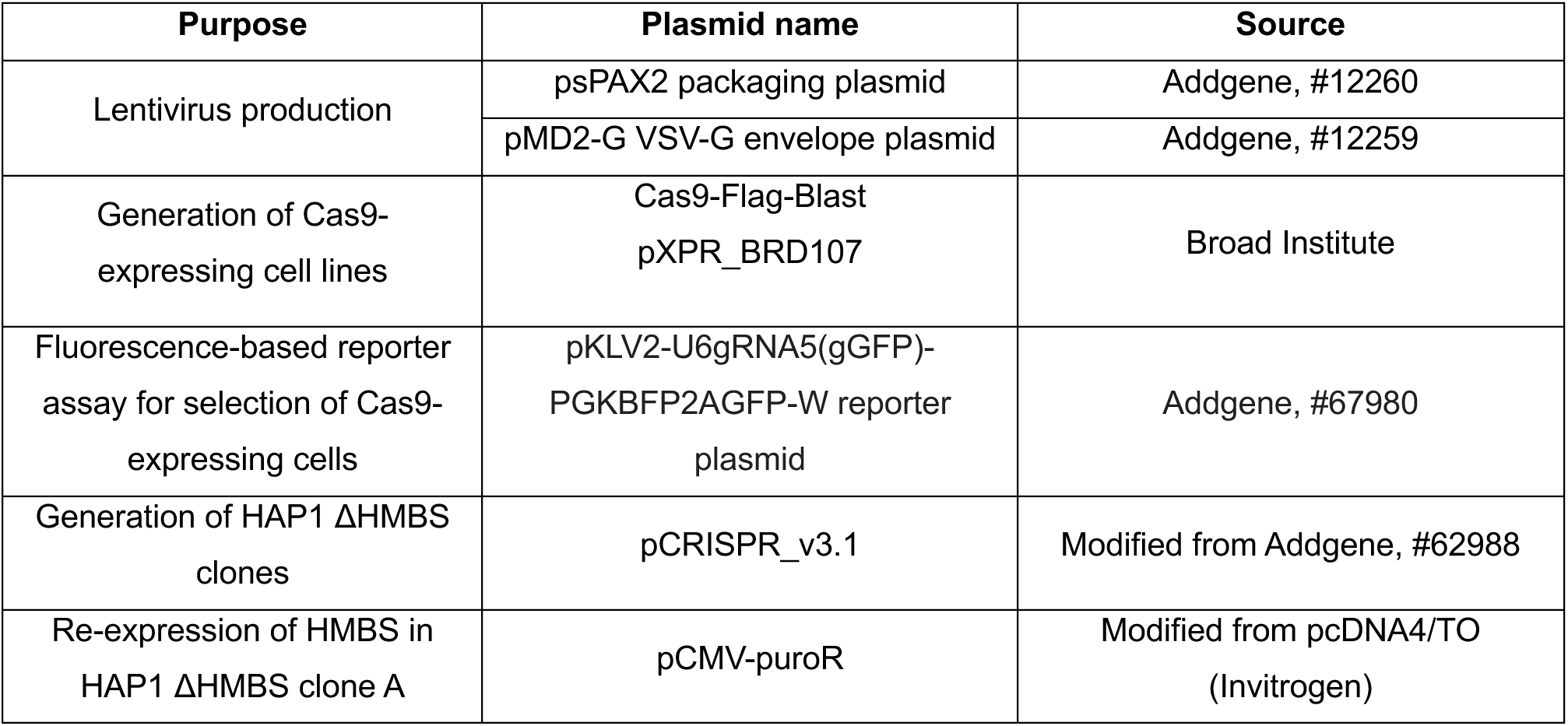
Plasmids employed in the study.

| Purpose | Plasmid name | Source |
| --- | --- | --- |
| Lentivirus production | psPAX2 packaging plasmid | Addgene, #12260 |
|  | pMD2-G VSV-G envelope plasmid | Addgene, #12259 |
| Generation of Cas9-expressing cell lines | Cas9-Flag-Blast<br>pXPR_BRD107 | Broad Institute |
| Fluorescence-based reporter assay for selection of Cas9-expressing cells | pKLV2-U6gRNA5(gGFP)-PGKBFP2AGFP-W reporter plasmid | Addgene, #67980 |
| Generation of HAP1 ΔHMBS clones | pCRISPR_v3.1 | Modified from Addgene, #62988 |
| Re-expression of HMBS in HAP1 ΔHMBS clone A | pCMV-puroR | Modified from pcDNA4/TO (Invitrogen) |

### Human and bovine sgRNAs

All sgRNAs used in this study are listed in the Table S7.

**Table S7.**
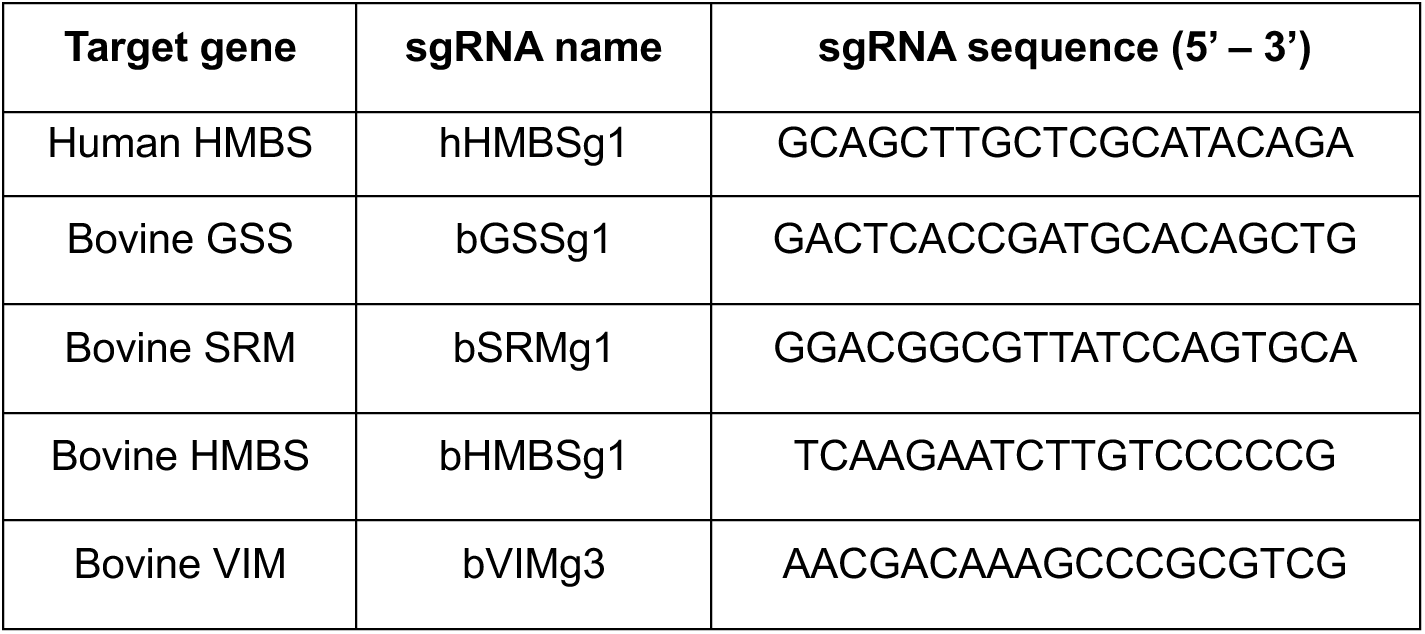
sgRNAs designed for single CRISPR/Cas9 knockout of candidate genes.

### Primers

All primers were synthesized by Microsynth AG (Baldach, Switzerland). Primers for plasmids and TIDE PCRs are listed in the Table S8.

**Table S8.**
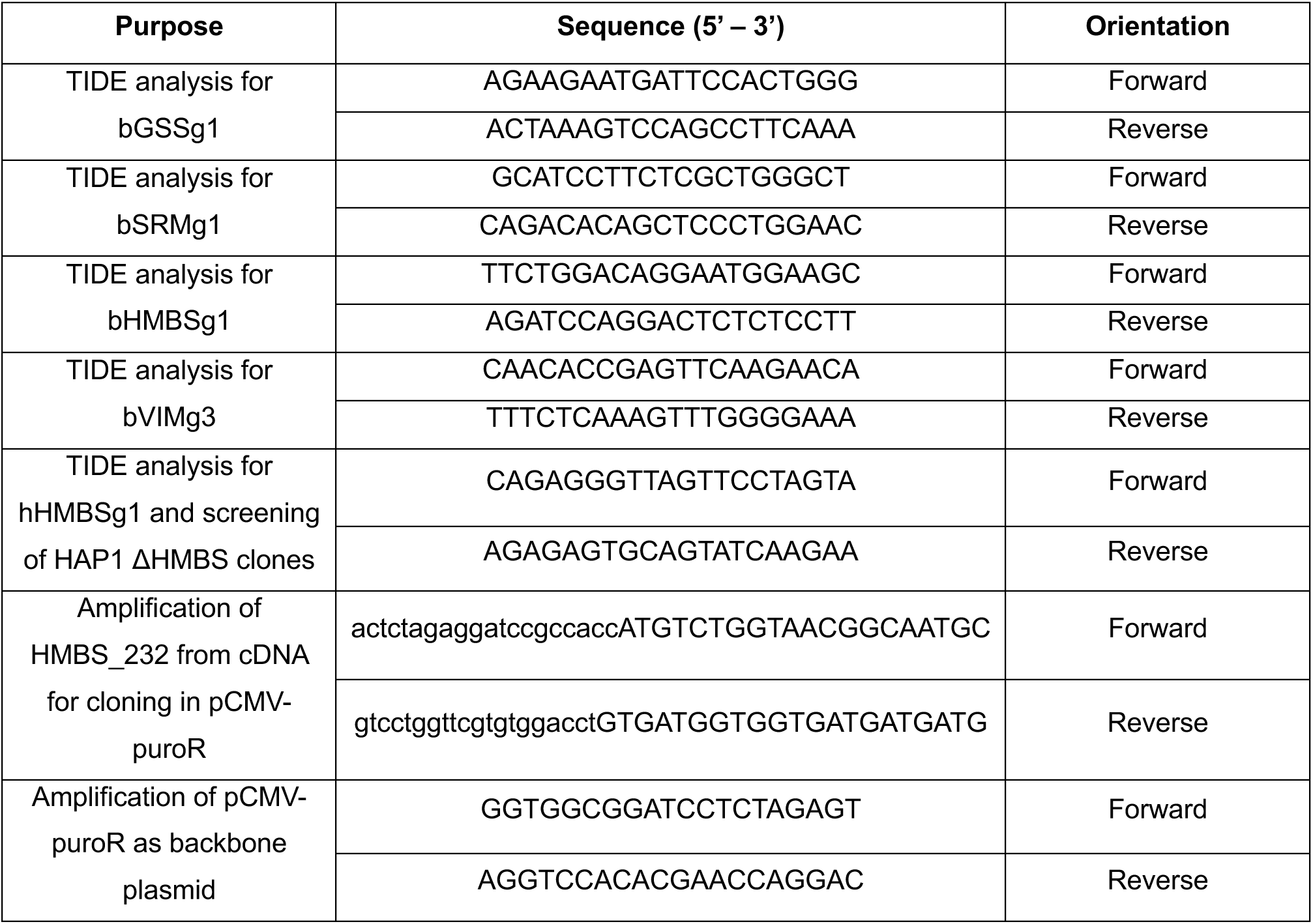
Sequences of primers. Lower-case letters indicate overhangs designed for cloning with NEBuilder HiFi DNA Assembly Master Mix.

### Lentivirus production

The following plasmids were used for lentivirus production: psPAX2 packaging plasmid (Addgene, #12260), pMD2-G VSV-G envelope plasmid (Addgene, #12259), Blast-Cas9 plasmid, pKLV2-U6gRNA5(gGFP)-PGKBFP2AGFP-W reporter plasmid (Addgene, #67980), and custom pooled CRISPR libraries. Lentiviruses were produced using the following protocol. First, ∼70% confluent HEK293FT cells in a 15 cm plate dish were transfected with transfer, packaging, and envelope vectors in a 4:2:1 ratio. 125 µl of 2.5 M calcium chloride was added to the plasmid mixture, and the final volume was made up to 1125 µl using 1 mM Tris, 0.1 mM EDTA, pH 8. The mixture was incubated at room temperature for 5 min. Then 1250 µl of 2 x HBS (50 mM Hepes, 280 mM NaCl) was added dropwise to the final plasmid solution while vortexing at full speed for 1 min. The mixture was immediately added dropwise to HEK293FT cells. Approximately 16 h post-transfection, the culture medium was replaced with fresh medium. The following day, ∼30 h after media replacement, cell supernatants were collected and filtered through a 0.45 µM filter to remove cell debris. Virus-containing supernatants were ultracentrifuged for 2 h at 20,000 x g at 4 °C (Hitachi CP100NX ultracentrifuge) and the pellet was aliquoted in 1 x PBS (∼200 µl for viruses produced from one 15 cm plate dish). Concentrated virus was aliquoted and stored at −80 °C. Viruses were harvested every 24 h (by adding fresh medium to the HEK293FT cells) for 3 - 4 days. The viral titer of concentrated viruses was calculated using the qPCR Lentivirus Titration Kit from ABM (Applied Biological Materials, Vancouver, Canada) according to the manufacturer’s instructions.

### Generation of Cas9-expressing cell lines

To achieve constitutive Cas9 expression, TaC12 and BoMac cell lines were infected with lentiviruses containing the Cas9-Flag-Blast construct at a multiplicity of infection of 10. Briefly, 100,000 – 500,000 cells were centrifuged at 100 x g for 5 min. The supernatant was removed, and the cell pellet was resuspended in 1 ml medium containing 8 µg/ml of polybrene. Aliquots of lentivirus stored at −80 °C were thawed on ice and the appropriate volume of virions was added to the cells. Virus-cell solutions were incubated for 20 min at room temperature, followed by a 30 min spin-infection performed at 30 °C in a laboratory centrifuge (100 x g). After centrifugation, the pellet was resuspended in the same virus-containing medium and cells were seeded into 6-well plates. After 24 h of incubation, the supernatant was replaced with fresh medium containing blasticidin (Sigma-Aldrich, St. Louis MO, USA) to select transduced cells (10 µg/ml for TaC12, 5 µg/ml for BoMac). For each cell line, two controls were included in the experiment: positive control (non-transduced cells in polybrene containing medium) and negative control (non-transduced cells in blasticidin containing medium). When the negative controls reached 100% cell death, transduced blasticidin-resistant cells were collected for validation of Cas9 expression by Western blotting. A fluorescence-based reporter assay was performed to specifically select cells expressing active Cas9. Briefly, Cas9-expressing cells were transduced with a lentiviral vector consisting of blue fluorescent protein (BFP) and green fluorescent protein (GFP) expression cassettes as indicated, together with a sgRNA targeting GFP. Cells were infected at an MOI of 10 in the presence of 8 µg/ml of polybrene for 24 h and seeded into T25 flasks. The culture medium was refreshed the following day. Between five and seven days after transduction, cells were sorted by FACS for BFP+GFP-signal, indicating expression of active Cas9 protein. For sorting, cells were resuspended in 1% FCS/PBS at a concentration of 10 mio cells/ml. BFP+GFP-cells were collected in a Falcon tube containing 100% FCS, centrifuged, and rapidly transferred to T25 flasks containing normal culture medium. Sorting was repeated twice for enrichment, using the MoFlo ASTRIOS EQ cell sorter (Beckman Coulter).

### CRISPR/Cas9 dropout screens

Genome-wide CRISPR/Cas9 dropout screens were performed in Cas9-expressing TaC12 cells using a two-vector system, as well as small-scale screens in TaC12 and BoMac cells. For the genome-scale screen we generated a CRISPR sgRNA containing 84,155 sgRNAs targeting 21,039 protein-coding genes, with 4 sgRNAs designed per each gene. For the small-scale screen we generated a CRISPR sgRNA library containing 16,596 sgRNAs targeting 1,561 protein-coding genes, with 10 sgRNAs designed per each gene. Five hundred non-targeting and 500 intergenic sgRNAs were added to each CRISPR library as controls. The bovine libraries were cloned into lentiGuide-Puro plasmid (Addgene, #52963) and viruses were produced in HEK293FT cells. Screens were performed at a coverage of 500 and in biological triplicates. A starting population of at least 140 mio cells (genome-wide screen) or 60 mio cells (small-scale screen) was infected with the bovine lentiviral library in normal growth medium in presence of polybrene (8 µg/ml) for 24 h. The next day, the culture medium was replaced and fresh medium containing puromycin (2 µg/mL for TaC12; 1 µg/mL for BoMac) was added to the cells. In both genome-wide and small-scale dropout screens, a positive control (untransduced cells) and a negative control (untransduced cells treated with puromycin) were included for each replicate of each CRISPR screen in all bovine cell lines. After 3 days of puromycin selection the negative control reached complete cell death, while successfully transduced cells were > 43 mio cells (e.g., 85,155 sgRNAs x 500) or > 9 mio cells (e.g., 17,596 sgRNAs x 500), corresponding to the previously calculated transduction efficiency of ∼30%. Puromycin-resistant cells were maintained in culture under drug selection for at least 11 passages and harvested at the first and last passages for genomic DNA (gDNA) extraction. gDNA was extracted from “START” and “END” point samples and high-throughput Illumina sequencing was performed to quantify the abundance of each sgRNA by PCR amplification of the sgRNA cassette. The resulting reads were deconvoluted using the PoolQ tool (https://portals.broadinstitute.org/gpp/public/software/poolq) to generate a matrix of read counts, and transformed into log-norm files. Log2 fold change (LFC) values for each guide were then generated by subtracting “END” point from “START” point values.

### Generation of HAP1 knockout cell line

For human HMBS knockout, we chose a sgRNA targeting an early coding region of the HMBS gene (5’-GCAGCTTGCTCGCATACAGA-3’) and cloned this guide into the pCRISPR_v3.1 plasmid. Low-passage HAP1 cells (< p5) were seeded at a concentration of 50,000 cells/well in 24 well plate. The following day, 600 ng of pCRISPR_v3.1+sgRNA (HMBS) plasmid was transfected using Lipofectamine3000 according to manufacturer’s instructions. Twenty-four hours post transfection, the culture medium was refreshed and 0.6 µg/ml puromycin was added to the transfected cells as well as in the negative control. Genomic DNA was extracted and the sgRNA (HMBS) target locus was PCR amplified using Q5 High-Fidelity DNA polymerase (New England Biolabs, Ipswich, Massachusetts) (forward primer: CAGAGGGTTAGTTCCTAGTA, reverse primer: AGAGAGTGCAGTATCAAGAA). The resulting amplicon was purified using the Gel & PCR clean-up kit (Macherey-Nagel, Düren, Germany) and sequenced by Sanger sequencing. The online tool Tracking of Indels by Decomposition (TIDE) was used to interpret the sequencing results. Limiting dilutions of the polyclonal HAP1 ΔHMBS pool were performed in 96 well plate, and six isolated clones were selected. Successful knockout of the HMBS gene was verified at genome level (identification of frameshift mutations at the target locus) and at the protein expression level (Western blotting).

### Re-expression of HMBS in HAP1 ΔHMBS clone A

The housekeeping isoform of HMBS transcript (ENST00000652429.1, HMBS_232) was amplified from complementary DNA synthesized by reverse transcriptase starting from RNA of HAP1 WT cells (GoScript^TM^ Reverse Transcriptase, Promega, Madison WI, USA). HMBS_232 was cloned into pCMV-puroR lentiviral plasmid using the primers listed in Table S6 and NEBuilder HiFi DNA Assembly Master Mix (New England BioLabs). Lentiviruses were produced as already mentioned. HAP1 ΔHMBS cells were transduced for 24h with virus-containing supernatants harvested from HEK293FT dishes, and cells were selected for 4 days with 0.6 µg/ml puromycin. Untransduced HAP1 ΔHMBS cells were used as a negative control and reached 100% cell death following 4 days of puromycin selection. Puromycin-resistant cells were subjected to limiting dilutions, and several single clones were analyzed via Western blotting for HMBS expression.

### Western blotting

Whole cell lysates were prepared by lysing 5 mio cells in 500 µl lysis buffer (50 mM Tris-HCl, 150 mM NaCl, 1 mM EDTA, 1% NP-40, 0.25% sodium deoxycholate, 1x cOmplete^TM^ protease inhibitor cocktail (Roche, Basel, Switzerland). The samples were boiled for 5 min at 90 °C, and then cooled on ice. Whole cell lysates were then loaded onto 10% sodium dodecyl sulphate (SDS) polyacrylamide gels. Gel runs lasted 45 - 50 min at 25 mA in electrophoresis buffer (2.5 mM Tris, 19.2 mM glycine, 0.01% SDS), followed by protein transfer to a nitrocellulose membrane for 90 min at 400 mA in transfer buffer (20 mM Tris-HCl, 15 mM glycine, 20% (v/v) methanol). The membrane was incubated for several hours in blocking solution (5% milk powder in TBST (10 mM Tris, 150 mM NaCl, 0.05% Tween). Primary antibodies were diluted in blocking solution and incubated with the membrane overnight at 4 °C. After three washes with TBS (10 mM Tris, 150 mM NaCl), the membrane was incubated with secondary antibodies diluted in blocking solution for 45 min at room temperature. If fluorescent secondary antibodies were used, incubation was performed in the dark. After three further washes with TBS, the Western blot was then imaged using the C600 imaging system (Azure Biosystems).

### FCS heme depletion

Heme-depleted FCS was prepared by treating FCS with 20 mM ascorbic acid for 16 h at 37 °C, followed by 24 h dialysis against PBS. Heme depletion was verified by measuring optical absorbance at 405 nm.^65^

### Cell viability assay

Cell viability assays were performed in 96 well plates. Briefly, 5,000 TaC12 or BoMac cells were seeded into the 96 wells and cells were allowed to grow for 72 h under normal culture conditions in the presence of inhibitors or solvent alone. Cell viability was measured using the resazurin assay.

### Mouse experiments and mosquito maintenance

Mouse studies were approved by the Commission for Animal Experimentation of the Cantonal Veterinary Office of Bern and conducted in compliance with the Swiss Animal Welfare legislation and under license BE118/22. Mice were infected by i.p. injection of blood stabilates of mCherry expressing *Plasmodium berghei* parasites (RMgm-928, 1804cl1). At approximately 4% parasitemia, blood was collected by cardiac puncture and passaged by i.v. injection into a new mouse (feed mouse). After 3 days, the parasitemia and gametocytemia were monitored by flow cytometry (NovoCyte, Agilent) and thin blood smear. At >0.4% gametocytes, the mice were anaesthetized (ketamin/xylazine) and made available to approximately 150 female *Anopheles stephensi* mosquitoes to take a blood meal for 30 min. The mosquitoes were maintained at 20.5 °C and above 80% relative humidity. Sporozoites were isolated from infected salivary glands on days 18 - 26 after the infective blood meal.

### Plasmodium berghei infection

HAP1 cells were cultured in IMDM (Bioconcept, containing 10% FCS, 4 mM L-glutamine, 100 U penicillin, 100 µg/ml streptomycin) at 37° C, 5% CO_2_. To infect confluent cultures, 70,000 cells were seeded in 96 well plates. The next day, the cultures were infected with 20,000 freshly isolated sporozoites. To allow development over 48 h, cultures were expanded by passaging with accutase (Innovative Cell Technologies) at 2 hpi. Cells were reseeded into 96 well plates (Greiner µClear, 655090). Each infected well was divided into 12 wells. At 48 hpi, cells were imaged with a Nikon Ti2 inverted fluorescence microscope (Plan Apo λD 10× (NA 0.45) objective, Spectra X light engine (555nm line, Lumencor), acquired with a Kinetix22 camera (Photometrix), Pinkel quad and Sedat quad filter sets (MXR00244 and MXR00254, Semrock). Parasite sizes were segmented and analyzed using the General Analysis Module of NIS-Elements 5.42.02. Graphs were generated with Prism 9 (GraphPad).

## SUPPLEMENTAL INFORMATION

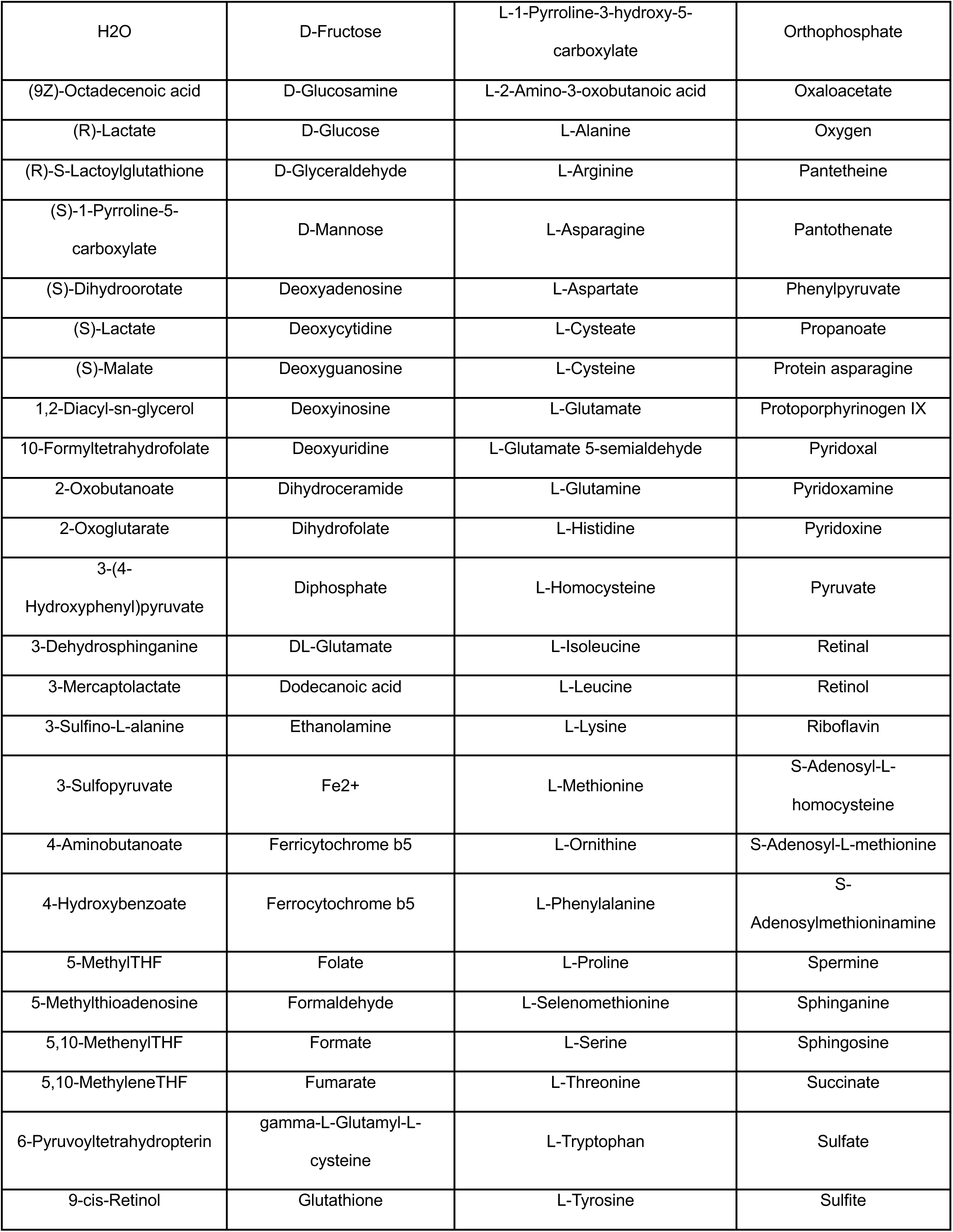
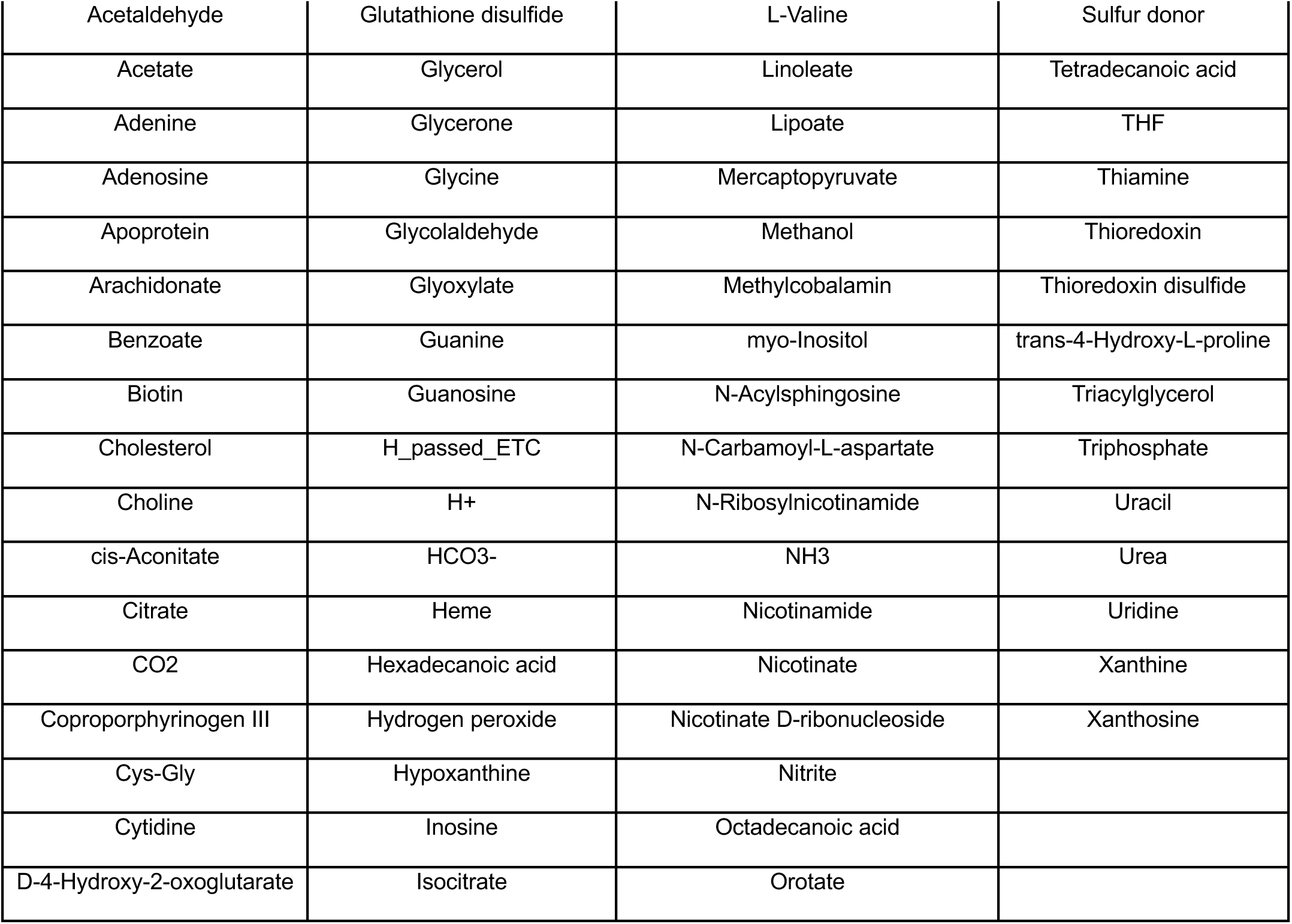

**Table S1. Composition of rich extra-parasitic environment.** 165 compounds that are available in the cytosol of the human Recon 3D model and that are part of the extracellular environment of the *Plasmodium falciparum* iPfa model. THF: tetrahydrofolate.

**Table S2. Hypergeometric analysis of bovine genome-wide CRISPR screen in TaC12 cells.** Average LFC values of the three screening replicates (Average_LFC) were normalized using the dispersion of intergenic and non-target controls (z-norm LFC).

**Table S3. Hypergeometric analysis of bovine small-scale CRISPR screen in TaC12 cells.** Average LFC values of the three screening replicates (i_Average_LFC; u_Average_LFC) were normalized using the dispersion of intergenic and non-target controls (i_z_LFC; u_z_LFC).

**Table S4. Fitness-scoring genes in small-scale CRISPR screen in TaC12 cells and dispensable for BoMac (*Theileria essentialome*)**. Average LFC values of the three screening replicates (i_Average_LFC; u_Average_LFC) were normalized using the dispersion of intergenic and non-target controls (i_z_LFC; u_z_LFC).

**Table S5. Gene Ontology analysis of *Theileria essentialome* (ShinyGO v.066).**

**Figure S1. Computational analysis of *P. falciparum* nutritional requirements and dependence on hepatocyte genes, related to Figure 2. (A)** Scheme of the computational workflow to generate *parasitosomes*: 1. Define the extra-parasitic environment; 2. Identify the reactions required for growth under the given conditions, and alternatives; 3. Extract the corresponding *parasitosome* reactions by collapsing the reactions identified in step 2 into one reaction containing the nutrients and products that each parasite state takes up and secretes into the host cytosol. **(B)** Composition of the compromised extra-parasitic environment, where the parasite has access to only the minimum set of metabolites (47 metabolites) required for growth. Participation in the alternative iMM sets determines whether each metabolite is constitutive, if it is required in all the alternatives, or substitutable, if another metabolite can substitute its function. **(C)** Metabolic subsystems composing the different networks for the *parasitosomes*, when considering a rich (green) or a compromised (orange) extra-parasitic environment (ePE), in terms of number of reactions and percentage of subsystem coverage. **(D)** Metabolic composition of the alternative *parasitosomes* in case study I (rich ePE, green) and II (compromised ePE, orange). The frequency indicates whether the nutrients and products are constitutive if they appear in all alternative *parasitosomes*.

**Figure S2. Computational analysis of *P. falciparum* nutritional requirements and dependence on hepatocyte genes, related to Figure 2**. Complete list of metabolic subsystems associated with hepatocyte genes that are essential for at least one *parasitosome*. Classification of the genes based on their frequency of essentiality across *parasitosomes*, the number of essential genes associated with each subsystem and the percentage of essential genes out of the total genes associated with each subsystem in case study I (rich ePE, green) and case study II (compromised ePE, orange).

**Figure S3. Hepatocyte genes essential for *P. falciparum* and dispensable for the host, related to Figure 2.** (**A**) List of genes essential in the healthy hepatocyte model. (**B - C**) Classification of genes per metabolic pathway and the frequency of their essentiality across *parasitosomes* in case study I (rich ePE, green) and in case study II (compromised ePE, orange). Genes essential for all *parasitosomes* are highlighted in red. (**D**) List of 19 genes essential for all *parasitosomes*.

**Figure S4. Cas9-expression in *Theileria*-infected macrophages and supplementary data on genetic screens, related to Figure 3. (A)** Western blot of whole cell lysates from Cas9-TaC12 and TaC12 WT. Flag-tagged Cas9 is expressed in TaC12 cells after transduction with Cas9-Flag-Blast lentiviruses. Tubulin staining served as a loading control. **(B)** Schematic representation of the lentiviral vector used for Cas9 activity assay. In transduced cells with no active Cas9 protein, both GFP and BFP fluorescence are detectable. When Cas9 is active, the GFP sequence gets cleaved and only the BFP fluorescence is detected. CMV, CMV promoter; RU5, 5’ long terminal repeat; hU6, human U6 promoter; gGFP, sgRNA targeting GFP gene; PGK, mouse *Pgk1* promoter; BFP, blue fluorescent protein gene; 2A, *Thosea asigna* virus 2A peptides; GFP, green fluorescent protein gene; W, Woodchuck hepatitis virus posttranscriptional regulatory element; ΔU3RU5, self-inactivating 3’ LTR. **(C)** TaC12-Cas9 expressing cells were analyzed by fluorescence-activated cell sorting (FACS) prior to cell sorting for BFP+/GFP-signal. The cell population expressing an active Cas9 protein (BFP+/GFP-) was sorted twice to enrich for BFP-positive cells. The BFP-positive gate was strictly set to enrich for the brightest fluorescing cells. Most cells transduced with the reporter construct are BFP+/GFP-while a small fraction is BFP/GFP double positive, indicating the absence of active Cas9. The same procedure was performed with BoMac-Cas9 cells. **(D)** Principal component analysis plot of log-norm count distribution of the TaC12 genome-wide screens. DAY0 and END points for each biological replicate are shown, as well as the distribution of the CRISPR library (log.library). **(E)** Library representation at DAY0 and END points of TaC12 genome-wide screen replicates. **(F)** Volcano plot showing the hypergeometric distribution analysis of the CRISPR/Cas9 small-scale screen in TaC12. Each gene is plotted based on its average LFC (z-normalized using the control dispersion) and the negative log_10_ of its *p*-value. A selected set of 100 genes that were found to be non-essential in TaC12_GW is shown in orange. A group of ∼ 800 genes significantly depleted in TaC12_GW (orig_TAC) is shown in yellow. The vertical dotted line corresponds to the average z-normalized LFC of the controls (intergenic and non-targeting) minus 1x standard deviation of the entire genetic screen. The screen was performed in 3 biological replicates. **(G)** Library representation at DAY0 and END points of TaC12 small scale screen replicates. **(H)** Volcano plot showing the hypergeometric distribution analysis of the CRISPR/Cas9 small-scale screen in BoMac. Each gene is plotted based on its average LFC (z-normalized using the control dispersion) and the negative log_10_ of its p-value. A selected set of 100 genes that were found to be non-essential in TaC12_GW is shown in orange. A group of ∼ 800 genes significantly depleted in TaC12_GW (orig_TAC) is shown in yellow. The vertical dotted line corresponds to the average z-normalized LFC of the controls (intergenic and non-targeting) minus 1x standard deviation of the entire genetic screen. The screen was performed in 3 biological replicates. **(I)** Library representation at DAY0 and END points of BoMac small scale screen replicates.

**Figure S5. TCA cycle genes are enriched in the *Theileria essentialome*, related to Figure 4. (A)** Representation of the TCA cycle by KEGG. Fitness-conferring genes in TaC12 are shown in red. **(B)** Scoring of genes belonging to the TCA cycle in TaC12 vs BoMac screens.

**Figure S6. Purine metabolism genes are enriched in the *Theileria essentialome*, related to Figure 4. (A)** Representation of purine metabolism by KEGG. Fitness-conferring genes in TaC12 are shown in red. **(B)** Scoring of genes belonging to the TCA cycle in TaC12 vs BoMac screens.

**Figure S7. Supplementary data related to Figure 5. (A)** TIDE analysis of *GSS*, *SRM* and *HMBS* CRISPR/Cas9 knockouts in TaC12 and BoMac cells. **(B)** TIDE analysis of vimentin (*VIM*) CRISPR/Cas9 knockout in TaC12 and BoMac cells. Vimentin is a non-essential gene in many cell lines, and we use it as a positive control that it can be knocked out in TaC12 and BoMac cells. **(C-E)** Resazurin viability assays showing the percentage of survival of BoMac and TaC12 cells after 72 hours treatment with buthionine sulfoximine (BSO), sodium nitroprusside (SNP), salicylic acid (SA). **(F)** Western blot of whole cell lysates from HAP1 WT and HAP1 ΔHMBS clones A-B-C-D-E-F. Tubulin staining served as loading control.

