## Supplementary figures and images for "Host metabolic pathways essential for malaria and related hemoparasites in the infection of nucleated cells"

### Supplemental Figures

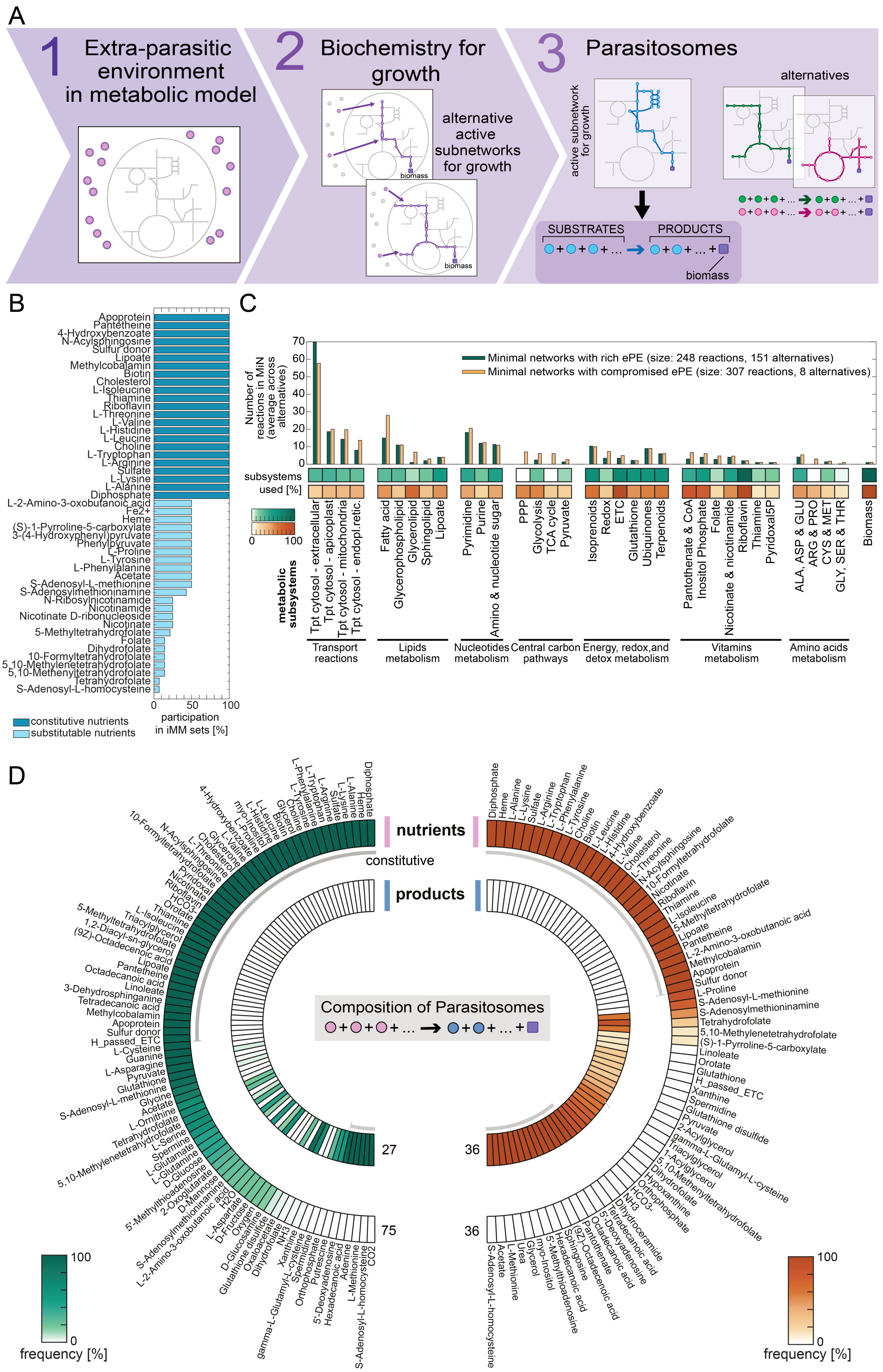

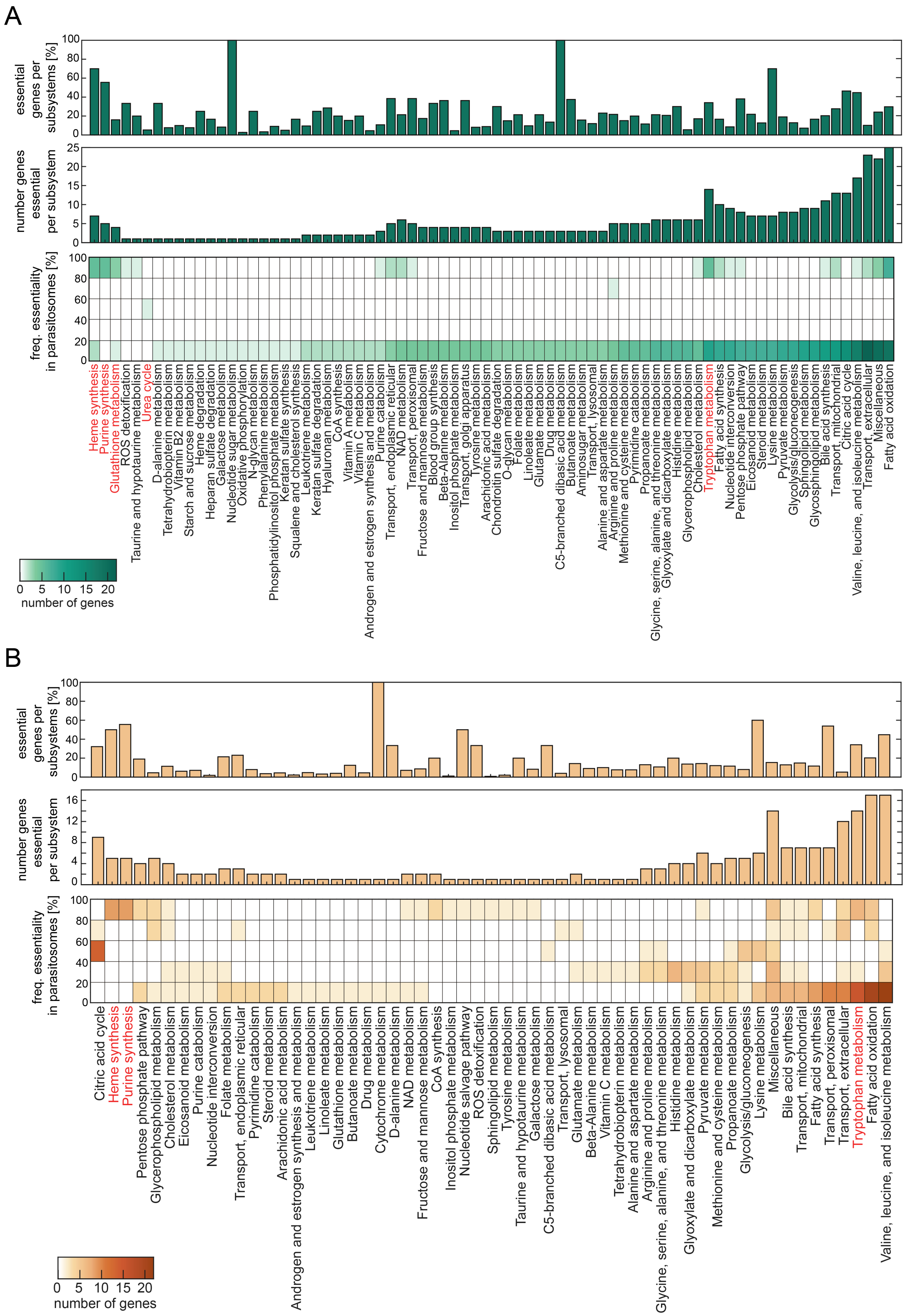

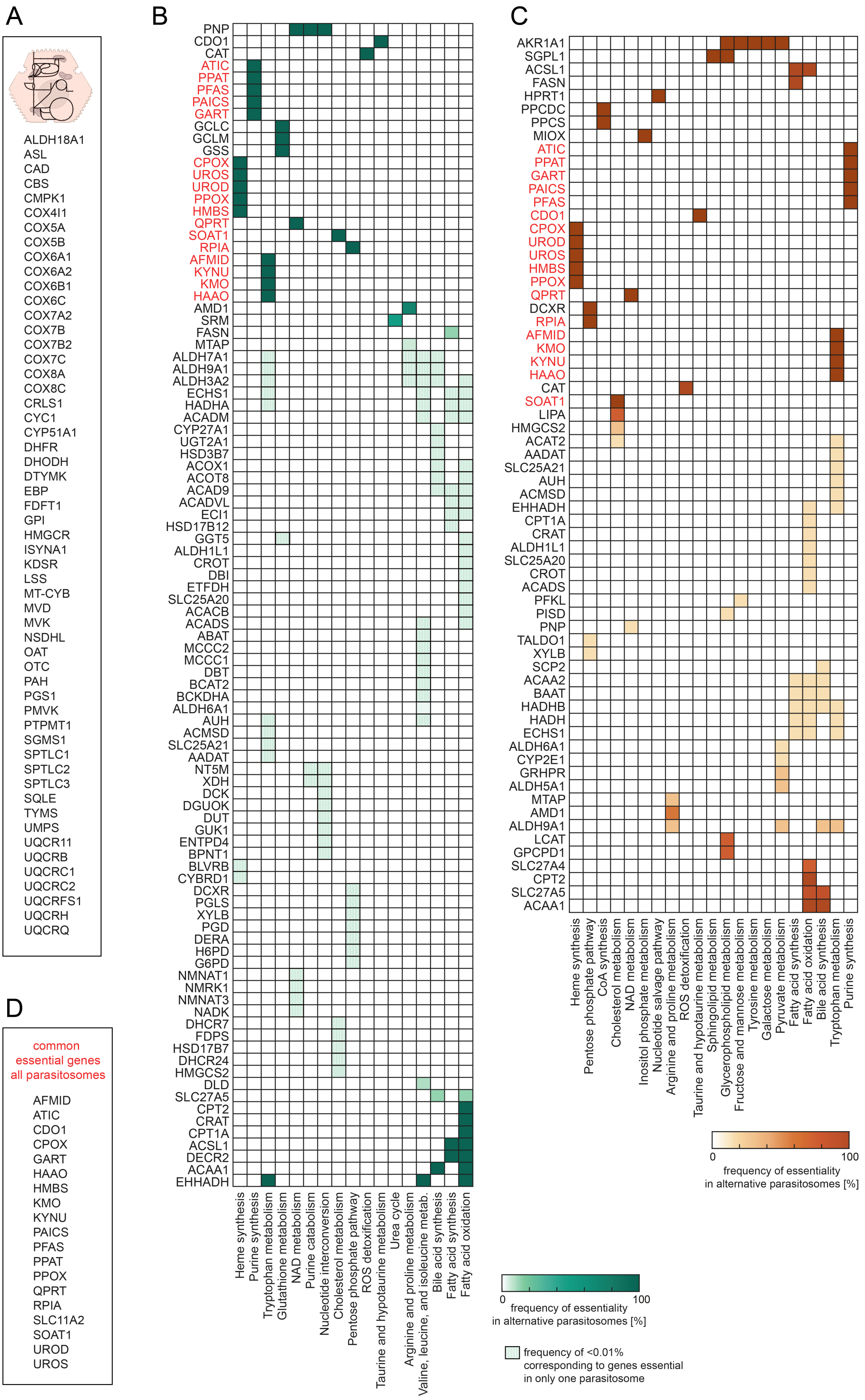

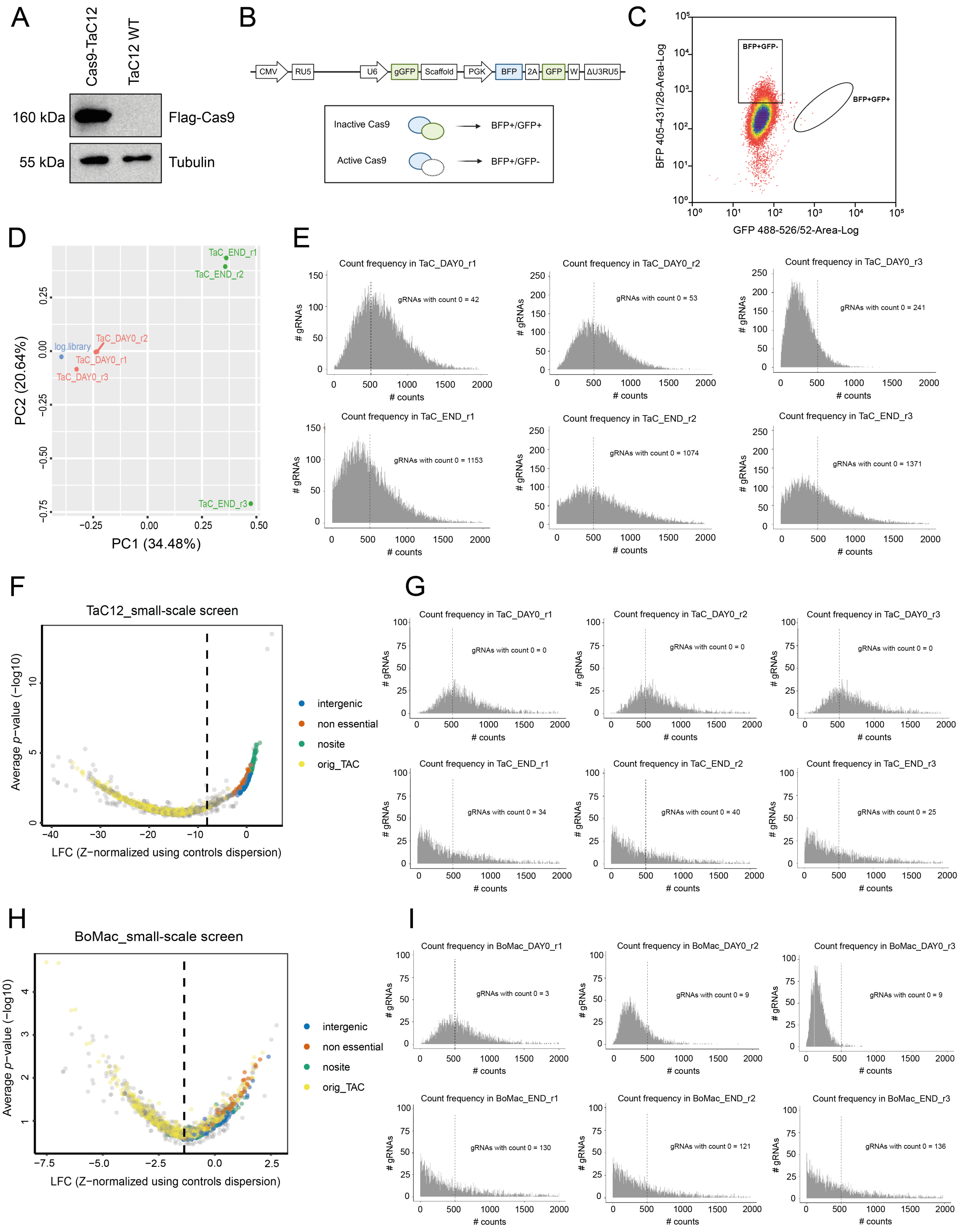

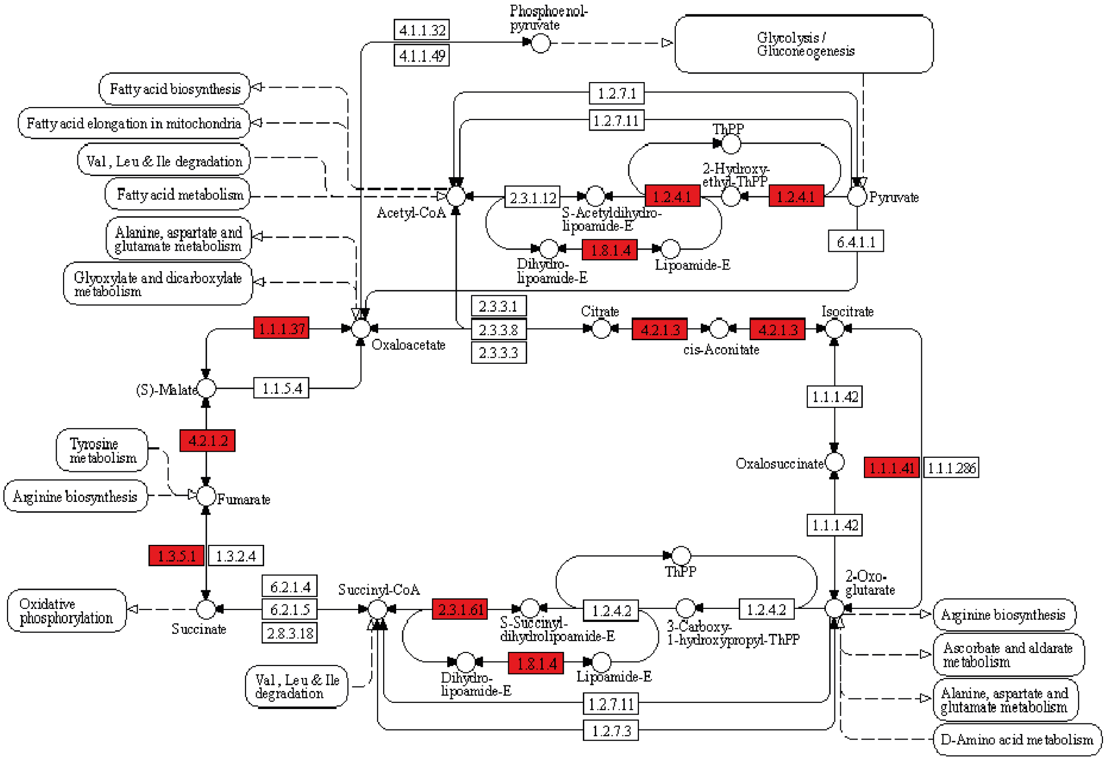

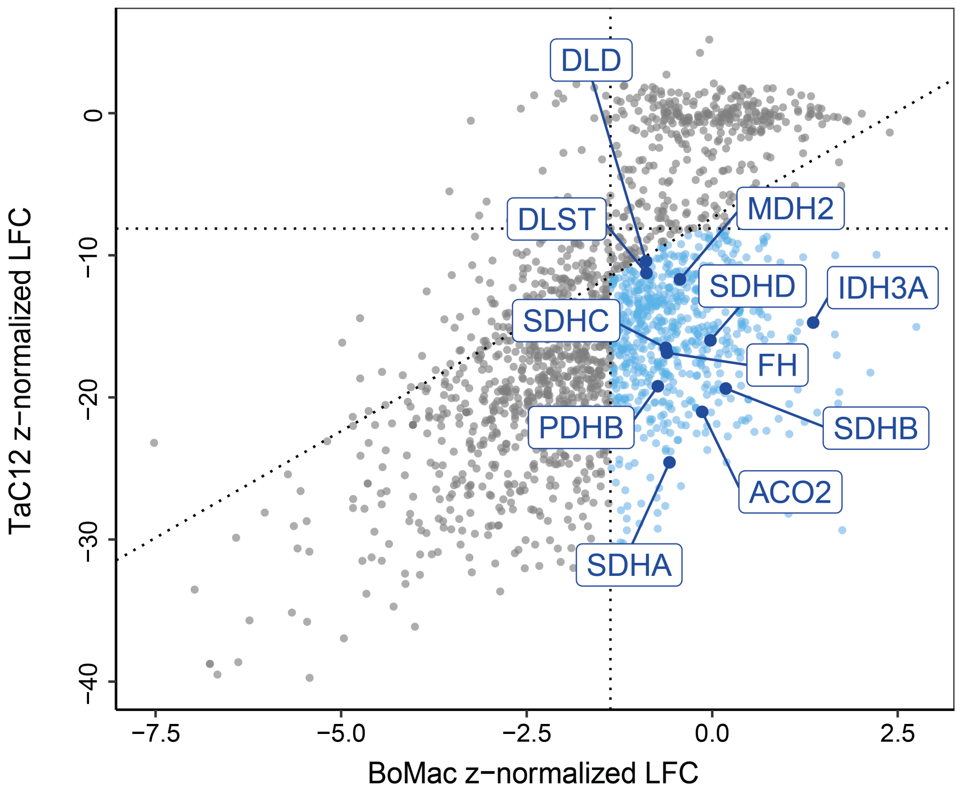

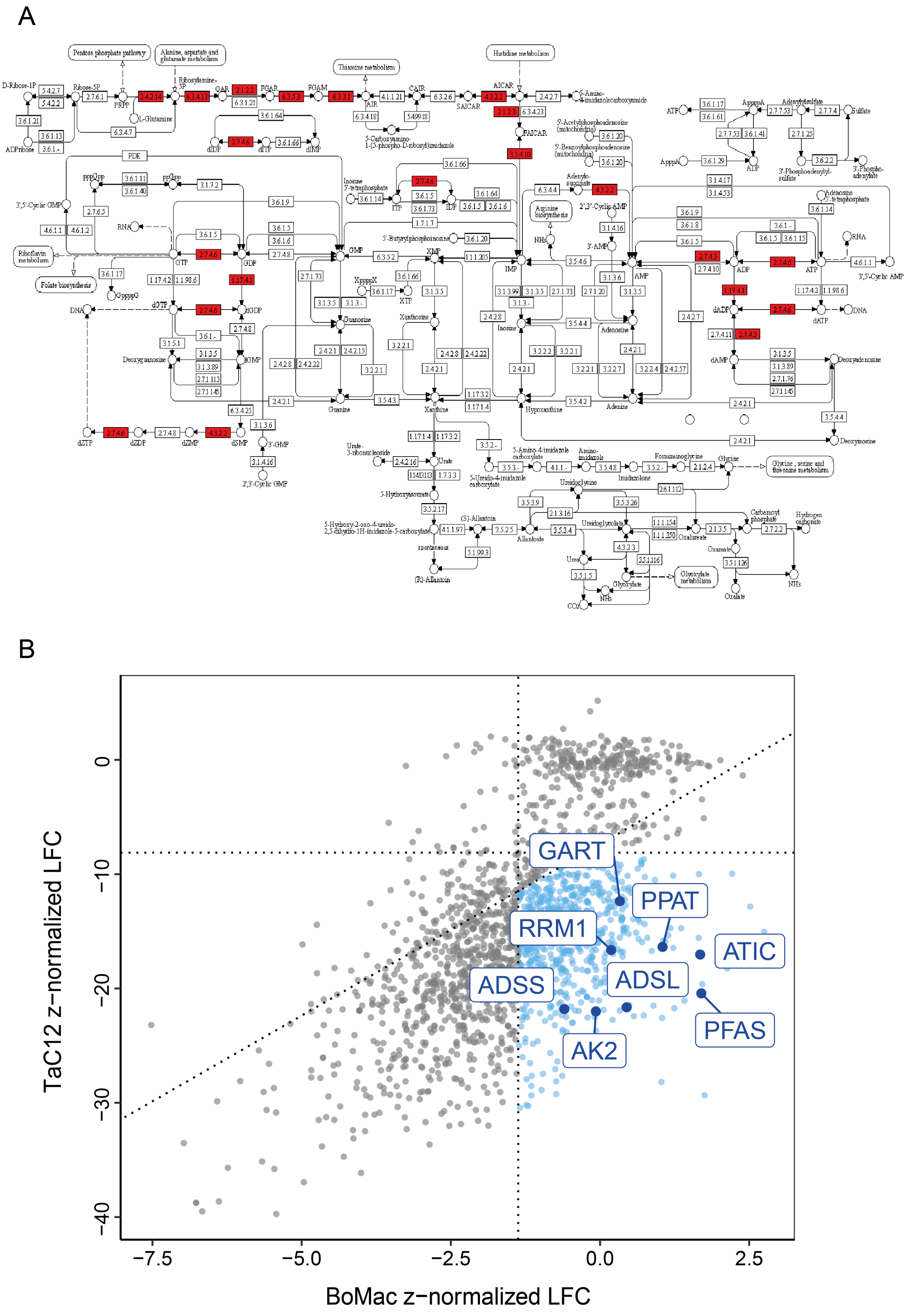

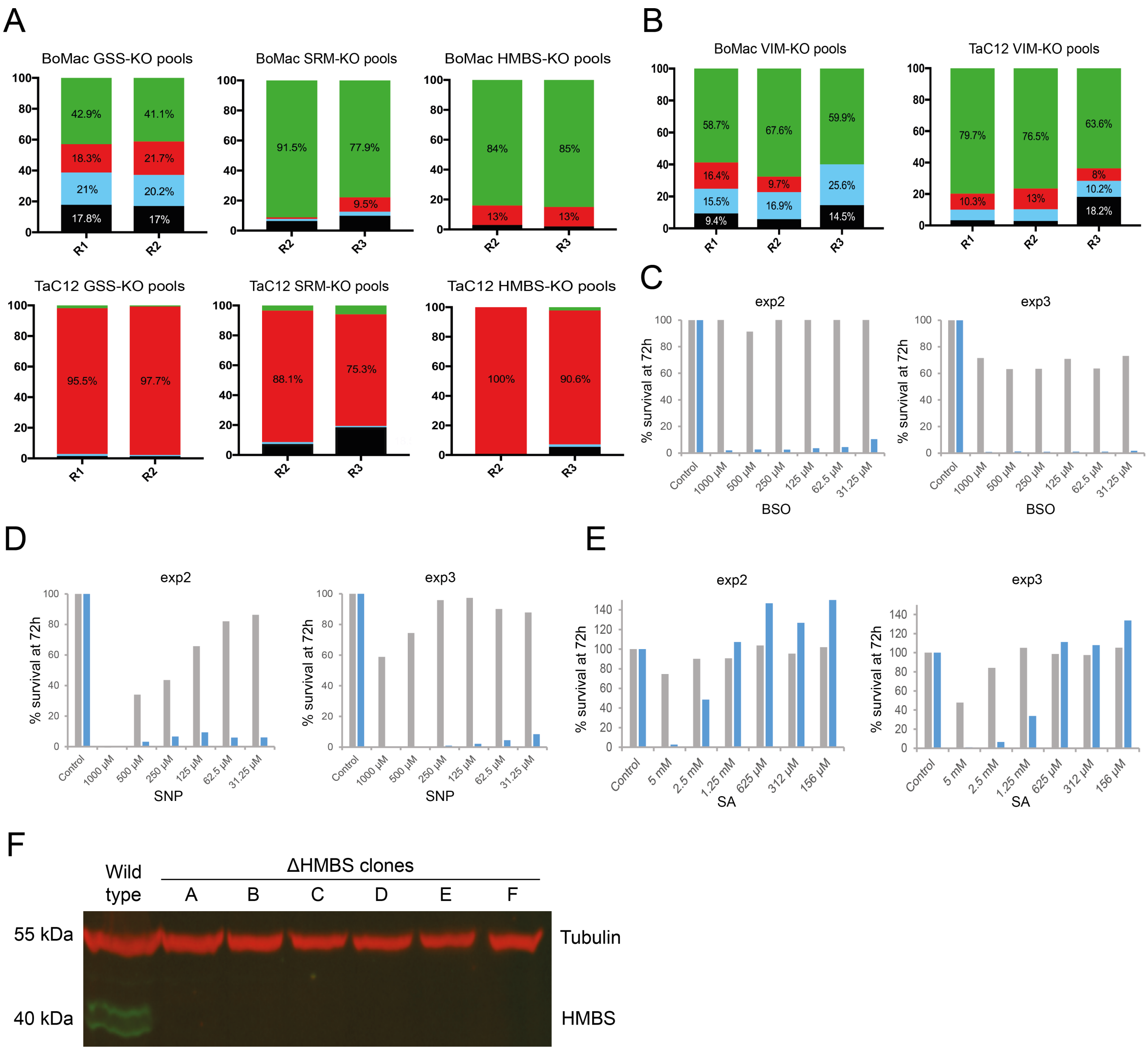
